# Lipid composition differentiates ferroptosis sensitivity between *in vitro* and *in vivo* systems

**DOI:** 10.1101/2024.11.14.622381

**Authors:** Vivian S. Park, Lauren E. Pope, Justin Ingram, Grace A. Alchemy, Julie Purkal, Eli Y. Andino-Frydman, Sha Jin, Sanjana Singh, Anlu Chen, Priya Narayanan, Sarah Kongpachith, Darren C. Phillips, Scott J. Dixon, Relja Popovic

## Abstract

Ferroptosis is a regulated non-apoptotic cell death process characterized by iron-dependent lipid peroxidation. This process has recently emerged as a promising approach for cancer therapy. Peroxidation of polyunsaturated fatty acid-containing phospholipids (PUFA-PLs) is necessary for the execution of ferroptosis. Ferroptosis is normally suppressed by glutathione peroxidase 4 (GPX4), which reduces lipid hydroperoxides to lipid alcohols. Some evidence indicates that GPX4 may be a useful target for drug development, yet factors that govern GPX4 inhibitor sensitivity *in vivo* are poorly understood. We find that pharmacological and genetic loss of *GPX4* function was sufficient to induce ferroptosis in multiple adherent (“2D”) cancer cell cultures. However, reducing GPX4 protein levels did not affect tumor xenograft growth when these cells were implanted in mice. Furthermore, sensitivity to GPX4 inhibition was markedly reduced when cells were cultured as spheroids (“3D”). Mechanistically, growth in 3D versus 2D conditions reduced the abundance of PUFA-PLs. 3D culture conditions upregulated the monounsaturated fatty acid (MUFA) biosynthetic gene stearoyl-CoA desaturase (SCD). SCD-derived MUFAs appear to protect against ferroptosis in 3D conditions by displacing PUFAs from phospholipids. Various structurally related long chain MUFAs can inhibit ferroptosis through this PUFA-displacement mechanism. These findings suggest that growth-condition-dependent lipidome remodeling is an important mechanism governing GPX4 inhibitor effects. This resistance mechanism may specifically limit GPX4 inhibitor effectiveness *in vivo*.

## INTRODUCTION

Ferroptosis is a non-apoptotic cell death mechanism that represents a potential target for cancer therapy^1,2^. This lethal mechanism is characterized by the iron-dependent accumulation of toxic membrane lipid hydroperoxides^3–7^. Membrane lipid peroxides accumulate on the endoplasmic reticulum, lysosome and potentially other organelles, and then at the plasma membrane^8^. Plasma membrane lipid peroxidation results in membrane stiffening and the opening of channels that alter the osmotic balance of the cell, ultimately resulting in cell lysis^9^. Understanding the factors that regulate sensitivity and resistance to ferroptosis is crucial to design therapies that target this mechanism.

The peroxidation of polyunsaturated fatty acid (PUFA)-containing phospholipids (PUFA-PLs) appears to be necessary for ferroptosis^10^. Several classes of PUFA-PLs are implicated in ferroptosis, including PUFA-containing phosphatidylethanolamines (PUFA-PEs), phosphatidylcholines (PUFA-PCs) and phosphatidylinositols (PUFA-PIs)^11^. Lethal lipid peroxidation can be inhibited by several mechanisms. The system x_c_^-^ cystine/glutamate antiporter, composed of solute carrier family 7 member 11 (SLC7A11) and solute carrier family 3 member 2 (SLC3A2), imports the anti-ferroptotic metabolite cystine into the cell in exchange for glutamate. Cysteine (the reduced form of cystine) can be used to synthesize (reduced) glutathione (GSH). GSH is synthesized in two steps by glutamate-cysteine ligase, which is comprised of a catalytic subunit (GCLC) and a regulatory subunit (GCLM), and glutathione synthetase (GSS). GSH is used by the selenoenzyme glutathione peroxidase 4 (GPX4) to reduce potentially toxic lipid hydroperoxides to lipid alcohols^10,12^. Additionally, lipid peroxidation reactions and ferroptosis can be inhibited by the ferroptosis suppressor protein 1 (FSP1)/coenzyme Q_10_/vitamin K^13–16^ and the GTP cyclohydrolase-1 (GCH1) and dihydrofolate reductase (DHFR)/dihydrobiopterin/tetrahydrobiopterin (BH2/4) pathways^17,18^. Despite these redundant anti-ferroptotic mechanisms, GPX4 inhibition alone is sufficient to induce ferroptosis in many cultured cancer cells. This has raised hopes for targeting GPX4 as an anti-cancer therapy. However, *Gpx4* is also essential for the survival of normal cells^19,20^. Toxicity associated with on-target GPX4 inhibition could therefore be problematic. Indeed, body weight loss is observed in mice following administration of GPX4 inhibitors^21,22^. For GPX4 to be a useful therapeutic target, a sufficient therapeutic window is needed so that ferroptosis can be induced in cancer cells, while normal cells are left unaffected. The factors that might increase or decrease the size of this window *in vivo* are poorly understood.

The ability of a cell to initiate ferroptosis is affected by the lipid composition of the membrane. The incorporation of less oxidizable monounsaturated fatty acids (MUFAs) in place of more oxidizable PUFAs into PLs, to yield MUFA-PLs, prevents ferroptosis *in vitro* and *in vivo*^5,23^. MUFAs can be taken up from the environment or synthesized *de novo* via stearoyl-CoA desaturase (SCD). MUFA free fatty acids are ‘activated’ by acyl-CoA synthetase long chain family member 3 (ACSL3) and incorporated into PLs by membrane bound O-acyltransferase domain containing 1/2 (MBOAT1/2)^5,23–29^. PUFAs and MUFAs compete for insertion into membrane PLs, with higher PUFA levels favoring ferroptosis execution and higher MUFA levels favoring ferroptosis inhibition^11^. Conditions that alter the balance of PUFA to MUFA species may impact our ability to induce ferroptosis to treat cancer and therefore require careful elucidation.

While investigating the regulation of GPX4 inhibitor sensitivity, we discovered that the same cancer cell lines were more sensitive to GPX4 inhibition when cultured in 2D adherent conditions than when cultured in non-adherent 3D (spheroid) conditions or in mouse models as tumor xenografts. Changes in the expression of key anti-ferroptotic genes did not sufficiently explain the increased resistance to ferroptosis in 3D environments compared to 2D. By contrast, alterations in the lipidome better explained observed differences in ferroptosis sensitivity between culture conditions. Culture in 3D conditions leads to upregulation of MUFA biosynthesis genes as well as enrichment of membrane phospholipids for MUFAs that promote ferroptosis resistance. Building on this insight, we identified specific MUFAs that protected against ferroptosis and pinpointed PUFA-PLs whose abundance was specifically impacted by MUFA metabolism. These results indicate that growth conditions can alter the lipidome and, by extension, GPX4 inhibitor sensitivities.

## RESULTS

### Ferroptosis sensitivity is altered by growth conditions

To develop effective ferroptosis-directed anti-cancer therapies^21,30^ we need to understand which targets in the pathway are essential for survival in different contexts. In cultured cells, ferroptosis is negatively regulated by several transporters and enzymes, including SLC7A11, GCLC/GCLM, GPX4, FSP1 and GCH1 (**Figure S1A).** To help guide our target selection and validation efforts, we examined the Broad Institute’s Dependency Map (DepMap)^31^ to determine which ferroptosis suppressor genes were most essential across cancer cell lines. Of 1,100 cell lines for which data were available, 66% were classified as dependent on GPX4 (732/1,100). By contrast, only 6.2% were dependent upon GCLC (68/1,100), 2% on SLC7A11 (22/1,100), 1.6% on GCH1 (18/1,100) and 1.1% on FSP1 (12/1,100) (**Figure S1A-C**). *GPX4* genetic dependency was not correlated with *GPX4* mRNA expression but was correlated with sensitivities to the small molecule GPX4 inhibitors 1*S*,3*R*-RSL3 (hereafter RSL3), ML162 and ML210, and to dependencies of other selenocysteine metabolic genes (e.g., *EFFSEC*, *PSTK*, *SEPSECS2*) (**Figure S1D-F**). This analysis suggested that many cancer cells are highly dependent upon GPX4.

Focusing on GPX4, we identified cancer cell lines that were sensitive (IC_50_ <1 µM) to RSL3-induced ferroptosis (**Figure 1A**). We initially selected for further examination 786-O clear cell renal cell carcinoma, Calu-1 non-small cell lung carcinoma (NSCLC) and NCI-H23 NSCLC as representative GPX4 inhibitor-sensitive models. In all three backgrounds we stably expressed Cas9 along with a CRISPR sgRNA targeting *GPX4* or a negative control region of the genome, under the control of a doxycycline (Dox)-inducible promoter. We refer to these cell lines as sg*GPX4* and sgControl (sgCTL), respectively (**Figure 1B**). GPX4 protein expression was reduced in all three Dox-treated (1 μg/mL, 72 h) sg*GPX4* cell lines (hereafter referred to as sg*GPX4^iKO^*) but not paired Dox-treated sgCTL cells (**Figure S2A**). sg*GPX4^iKO^* cells had lower cell viability, as measured using CellTiter-Glo, compared to sg*GPX4* cells treated with vehicle (DMSO) or sgCTL cells treated with DMSO or Dox (**Figure 1B**). These results were obtained using cells growing in 2D cultures. Using these three cell models, we investigated whether GPX4 genetic disruption altered tumor growth *in vivo*. All three cell models were implanted as xenografts in severe combined immunodeficiency disease (SCID) and SCID-Beige mice fed a Dox-containing diet for seven days prior to tumor implantation. Once tumors reached a group median of 1000 mm^3^ or 110 days, tissue was harvested for downstream analyses. Surprisingly, no reduction in tumor growth was observed in any model despite GPX4 protein abundance being reduced by 90-95% in 786-O and Calu-1 sg*GPX4^iKO^* tumors, and 65-80% in NCI-H23 sg*GPX4^iKO^* tumors (**Figure 1C**).

**Figure 1.**
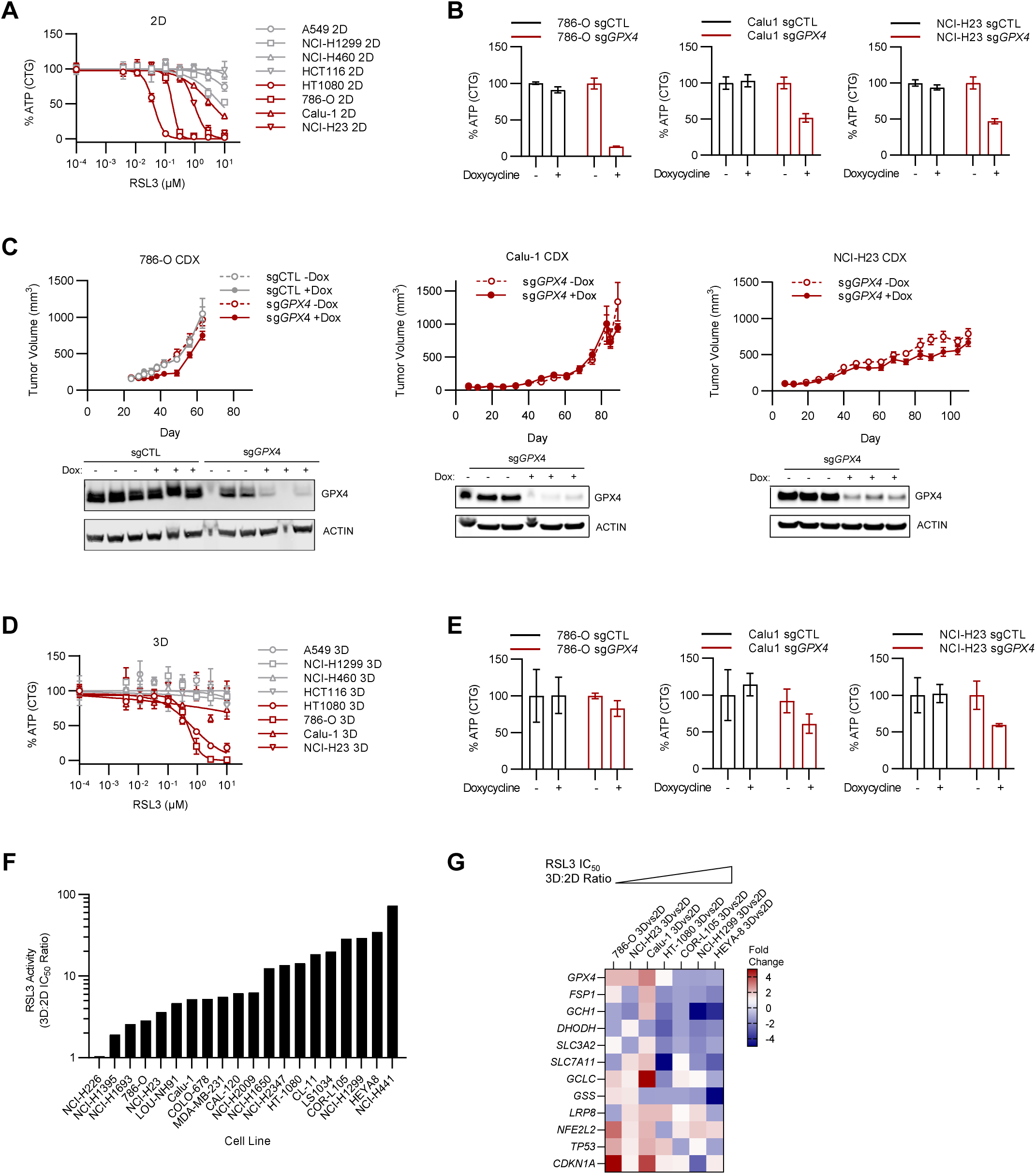
Cancer cell line ferroptosis sensitivity in 2D, 3D cells and *in vivo* tumors. **A.** ATP levels measured by CellTiter-Glo (CTG) for cancer cell lines treated with RSL3 in 2D cell culture. **B.** ATP levels (CTG) of inducible *GPX4* CRISPR/Cas9 knockout (KO) and non-targeting control (NTC) cell lines treated with DMSO or doxycycline (1μg/mL) in 2D cell culture. **C.** Tumor growth curve of inducible sg*GPX4^iKO^* cell-line derived xenograft (CDX) murine models. 786-O: sgCTL, sgCTL+Dox, sg*GPX4*, sg*GPX4*+Dox (n=10 for each treatment arm). Calu-1: sg*GPX4*, sg*GPX4*+Dox (n=10 for each treatment arm). NCI-H23: sg*GPX4*, sg*GPX4*+Dox (n=10 for each treatment arm). Western blot of GPX4 and Actin at endpoint, n=3 for each treatment arm. **D.** ATP levels (CTG) of cancer cell lines treated with RSL3 in 3D cell culture. **E.** ATP levels (CTG) of inducible *GPX4* KO cell lines treated with DMSO or doxycycline (1mg/mL) in 3D cell culture. **F.** RSL3 IC_50_ 3D:2D ratios of indicated cell lines. **G.** Heatmap of baseline expression changes (qPCR) of ferroptosis-related genes in 3D versus 2D for indicated cell line. Median of 3 biological replicates is represented. 2D cell culture of given cell line set as reference (=1).

It seemed that sg*GPX4^iKO^* xenograft tumor growth was unaffected, despite substantial loss of GPX4 protein. One possibility was that the *in vivo* tumor microenvironment suppressed ferroptosis. Another possibility was that growth in non-adherent, 3D conditions was sufficient to reduce ferroptosis sensitivity. To investigate the latter possibility, we cultured cells as spheroids (i.e., 3D culture) using ultralow attachment (ULA) plates, allowing spheroid formation for 48 h before downstream assays. Crucially, 3D and 2D cultures both used the same culture medium, so that differences in nutrient availability or growth factors should not contribute to the observed phenotypes. RSL3 was less effective at reducing cell viability in HT-1080, NCI-H1299, Calu-1 and NCI-H23 cells than when these same cells were cultured in 2D conditions (**Figure 1D** versus **1A**). Likewise, sg*GPX4^iKO^* cells had higher levels of viability in 3D conditions, although Calu-1 and NCI-H23 sg*GPX4^iKO^* 3D cells did maintain some sensitivity to cell death (**Figure 1E**). To extend these results further, we examined cell death responses in 18 normally adherent cancer cell lines from various lineages (lung, renal, breast and colorectal) in 3D and 2D conditions. Strikingly, the RSL3 IC_50_ was on average 14.5-fold (range: 1.1-fold to 73-fold) higher in 3D versus 2D culture (**Figure 1F**). For example, the sensitivity of HT-1080 and NCI-H1299 cells to RSL3 was reduced approximately 14-fold and 29-fold, respectively, in 3D versus 2D cultures (**Figure 1F**). While the 3D:2D RSL3 IC_50_ ratios varied across cell lines, greater resistance to RSL3 was observed in 17/18 cell lines cultured in 3D conditions compared to 2D. Thus, 3D growth conditions generally suppressed ferroptosis sensitivity in the cell lines examined here.

We hypothesized that enhanced ferroptosis resistance in 3D versus 2D growth conditions was explained by transcriptional upregulation of canonical ferroptosis suppressor genes lying in the *GPX4*, *AIFM2*/*FSP1*, and other known pathways. To test this hypothesis, we measured the expression of 12 ferroptosis suppressor genes in a panel of cell lines in both 3D and 2D cultures using high-throughput qPCR. Across all cell lines, differences in mRNA levels between 3D and 2D cultures did not clearly associate with ferroptosis resistance (**Figure 1G**). In fact, cell lines with higher 3D:2D RSL3 IC_50_ ratios sometimes expressed lower levels of ferroptosis resistance genes (e.g., *AIFM2/FSP1*, *GCH1* and *SLC7A11*) in 3D cultures versus 2D cultures. This suggested that other factors were responsible for the differences in ferroptosis sensitivities in 3D versus 2D conditions.

We next hypothesized that enhanced ferroptosis resistance in 3D versus 2D growth conditions was explained by better adaptation to GPX4 inhibition. For example, a ferroptosis resistance gene could be specifically upregulated in 3D versus 2D cultures in response to GPX4 inhibition. To investigate, we used genome-wide microarray data to analyze the transcriptomes of HT-1080 and NCI-H1299 cells cultured in 3D or 2D conditions and treated with DMSO (vehicle control) or RSL3 (0.1 μM, 2 h). RSL3 treatment of 2D HT-1080 or 2D NCI-H1299 cells did not significantly alter the expression of any genes (FDR *P* < 0.05) (**Figure S2B**). By contrast, treatment of 3D HT-1080 cells with RSL3 increased the expression of 126 genes and decreased the expression of 67 genes (>2 fold, FDR *P* < 0.05) compared to DMSO control. RSL3-treated 3D NCI-H1299 cells increased the expression of 7,527 genes and decreased expression of five genes. Interestingly, *HMOX1* and *SLC7A11*, two known regulators of ferroptosis^32,33^, were markedly upregulated in both 3D HT-1080 and 3D NCI-H1299 cells. (**Figure S2C**).

To investigate whether any single gene was functionally responsible for increased ferroptosis resistance in 3D growth conditions, we performed genome-wide CRISPR screens in NCI-H1299 cells^34^. First, we compared 3D versus 2D cell culture essentiality in NCI-H1299 cells. Consistent with a successful screen, genes known to be essential for survival in 3D conditions (e.g., *CPD* and *IRS1*)^35^ were depleted in 3D cells compared to 2D cells (**Figure S2D**). Next, we performed a genome-wide screen in 3D NCI-H1299 cells treated with RSL3 versus vehicle control (DMSO). However, comparison of RSL3-treated versus DMSO-treated 3D cells did not identify any single gene dependencies (FDR *P* < 0.05) (**Figure S2E**). Neither *SLC7A11* nor *HMOX1* emerged as depleted genes in our 3D screen. Indeed, even though *SLC7A11* expression was significantly upregulated in RSL3-treated 3D culture (**Figure S2C**) it might not be an expected dependency since it acts upstream of GPX4. Together, these data indicated that the increased resistance to chemical or genetic induction of ferroptosis in 3D cell culture and *in vivo* tumors was not explained by transcriptional differences of canonical ferroptosis negative regulators or any single genetic perturbation.

### Growth conditions alter lipid metabolism

Lipid metabolism is highly sensitive to growth conditions^36,37^. We therefore hypothesized that changes to the lipidome in 3D cultures and *in vivo* versus in 2D cultures could explain reduced ferroptosis sensitivity. To test this hypothesis, we compared the lipidomes of 786-O, Calu-1 and NCI-H23 cell lines growing as tumor xenografts, and in 3D or 2D culture conditions (**Figure 2A**). Principal component analysis of lipid classes in 786-O cells clustered based on growth conditions (i.e., *in vivo* tumors, 3D culture and 2D culture) (**Figure 2B**). Comparing the percent distribution of lipid classes in 786-O, Calu-1, and NCI-H23 cell lines, all three showed reduced phosphatidylcholine (PC), phosphatidylethanolamine (PE), and phosphatidylserine (PS) phospholipid abundance and a corresponding increase in triglyceride (TG) abundance when grown *in vivo* or in 3D compared to 2D (**Figure 2C**).

**Figure 2.**
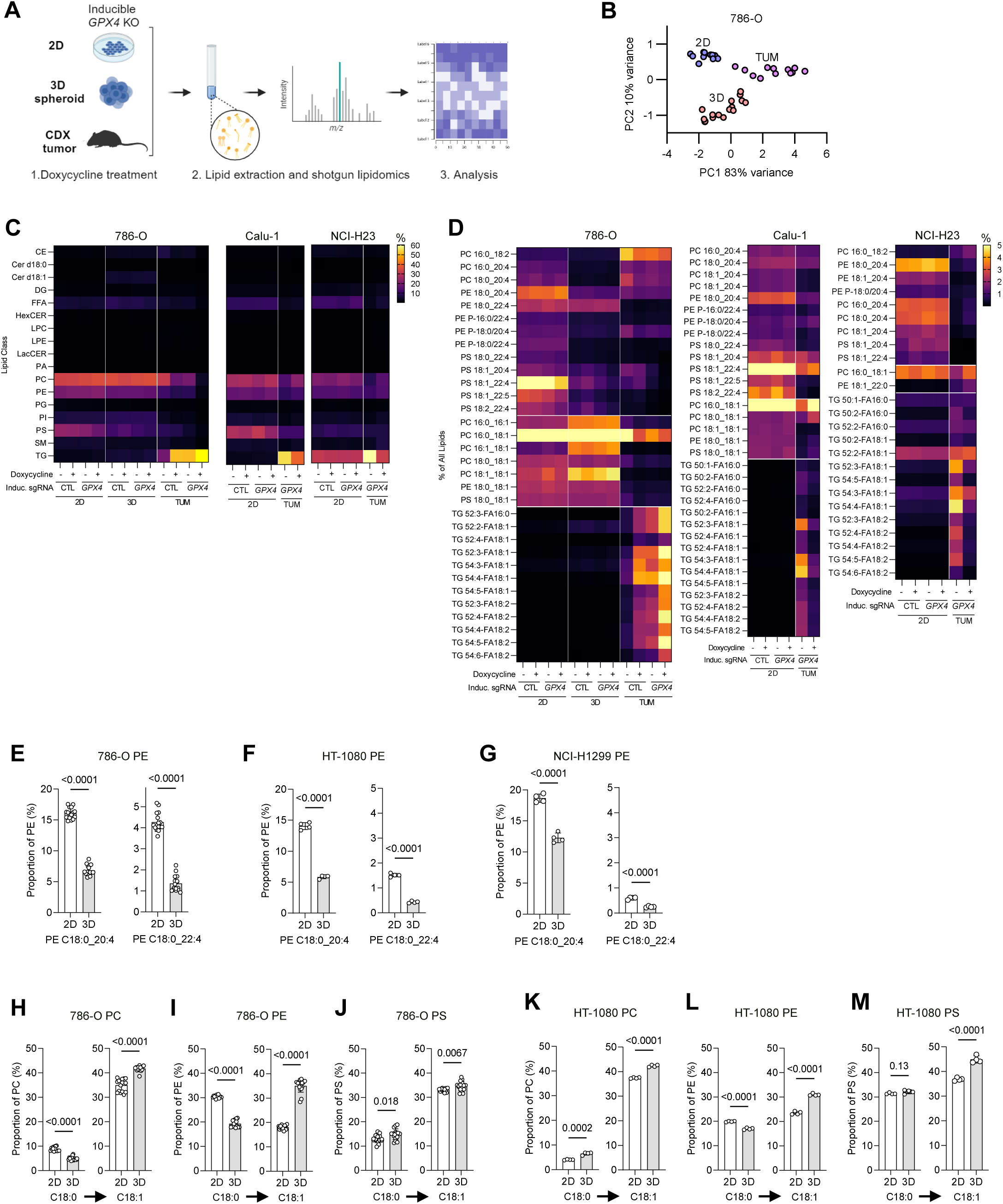
Lipidomic analyses of inducible *GPX4* KO cell line in different conditions. **A.** Schematic of the lipidomic analyses. Created with BioRender.com. **B.** Principal component analysis (PCA) of lipid classes in 786-O inducible *GPX4* KO cell line. Each replicate for 2D cells, 3D cells and tumors represented. **C.** Heatmap of proportion of each lipid class compared to total lipidome. For 2D and 3D cells, the median of 4 biological replicates is represented. For tumors, median of 3 biological replicates is represented. **D.** Heatmap of proportion of each lipid species compared to total lipidome. For 2D and 3D cells, the median of 4 biological replicates is represented. For tumors, median of 3 biological replicates is represented. **E**-**G.** Proportion of indicated lipid species PE-AA, PE-AdA, within PEs comparing 2D cells and 3D cells. **H**-**J** and **K**-**M.** Changes in 18:0-PL (saturated fatty acid, SFA) ◊ 18:1-PL (oleic acid, OA) within PC (**H**, **K**), PE (**I**, **L**) and PS (**J**, **M**) in 2D cells and 3D cells of 786-O inducible *GPX4* KO and HT-1080 cell lines. In **E**-**M**, data are presented as means ± SD and statistically significant differences were determined using two-tailed unpaired Student’s t-tests. *Abbreviations:* arachidonic acid (AA), adrenic acid (AdA), saturated fatty acid (SFA), oleic acid (OA), cholesterol esters (CE), ceramides (Cer d18:1), diacylglycerol (DG), dihydroceramides (Cer d18:0), free fatty acid (FFA), hexosylceramides (HexCER), lactosylceramides (LacCER), lysophoshpatidylcholine (LPC), lysophosphatidylethanolamine (LPE), phosphatidic acid (PA), phosphatidylcholine (PC), phosphatidylethanolamine (PE), phosphatidylglycerol (PG), phosphatidylinositol (PI), phosphatidylserine (PS), sphingomyelin (SM), triglyceride (TG).

Further analysis of lipid species revealed changes in lipid composition across experimental systems (**Figure 2D**). Lipid species were categorized based on the number of carbon-carbon double bonds as PUFA (two or more C=C double bonds) or MUFA (one C=C double bond). We then compared the proportions of lipid species (>2% compared to all lipids) between systems. Changes in overall PUFA content between growth conditions across all three models were observed. 786-O cells cultured as 3D spheroids or xenograft tumors both had fewer PUFA-containing species compared to cells grown in 2D. Likewise, Calu-1 and NCI-H23 cells had fewer PUFA-containing species when grown as xenografts compared to 2D cultures (**Figure 2D**). Of note, in 786-O, Calu-1 and NCI-H23 sg*GPX4* cell lines, Dox treatment (sg*GPX4^iKO^*) did not affect these patterns in 2D versus 3D cultures; however, sg*GPX4^iKO^* tumor cells had shifts in lipid composition. TG abundance in sg*GPX4^iKO^* tumors was markedly different compared to controls. Specifically, 786-O sg*GPX4^iKO^* tumors were enriched in TG lipid species compared to its controls. Calu-1 sg*GPX4^iKO^*and NCI-H23 sg*GPX4^iKO^* tumor cells, on the other hand, harbored fewer TG lipid species compared to their respective controls.

TGs can regulate ferroptosis by sequestering more oxidizable PUFAs away from PLs^11^. Given the observed enrichment for TGs in 786-O, Calu-1 and NCI-H23 xenograft tumors compared to cultured cells (**Figure 2D**), we examined the proportion of fatty acids within TGs, categorized by the number of double bonds. Notably, in 786-O tumors overall PUFA-TG content was higher compared to 2D and 3D cultures; tumor TGs were comprised of 20-30% C18:2 (likely linoleic acid) compared to 3-4% C18:2 in both 3D and 2D cultures. Enrichment of C18:2 was also observed in Calu-1 and NCI-H23 tumors versus respective 2D cultures (**Figure S3A**). In fact, C18:2 represented 60-80% of all tumor PUFA-TGs in the three models, whereas PUFA-TGs with three or more C=C double bonds were relatively diminished. This suggested that changes in TG MUFA:PUFA ratios were unlikely to account for the observed differences in ferroptosis sensitivity between xenograft tumors and *in vitro* culture systems. Furthermore, given the central importance of PUFA-containing PLs for ferroptosis execution, we focused mainly on this lipid class.

Next, we examined individual PUFA-containing and MUFA-containing PL species. We specifically focused on PUFA-PLs classically associated with the execution of ferroptosis: PEs containing arachidonic acid (AA, C20:4) and adrenic acid (AdA, C22:4)^3,38^. PE C18:0_C20:4 (PE-AA) was significantly less abundant in HT-1080, 786-O and NCI-H1299 3D versus 2D cultures (**Figure 2E-G**). PE-AA was also less abundant in 786-O, Calu-1 and NCI-H23 xenograft tumors compared to 2D cell cultures (**Figure S3B, SC**). PE C18:0_C22:4 (PE-AdA) was likewise significantly less abundant in HT-1080, 786-O, and NCI-H1299 3D versus 2D cultures (**Figure 2E-G**). Lower PE-AdA levels were also observed for 786-O and Calu-1 cells xenograft tumors versus 2D cultures (**Figure S3B, S3C**). Focusing on MUFA-PLs, 786-O cells were enriched for MUFA-PLs containing C18:1 in 3D versus 2D cultures (**Figure 2D**). Our analysis method did not allow us to resolve which specific C18:1 MUFA(s) were present in our lipids. However, one common C18:1 MUFA, *cis*-9-C18:1 (oleic acid, OA), is synthesized from the precursor saturated fatty acid (SFA) stearic acid (C18:0)^39^. Consistently, the proportion of C18:0 for 786-O, HT-1080 and NCI-H1299 PEs was reduced in 3D versus 2D cultures (**Figure 2I, 2L, S3E**).

Increased MUFA levels can promote a ferroptosis-resistant state^5,23,24,27,40,41^. We therefore once again abstracted the results to a higher level to compare overall MUFA-PL versus PUFA-PL content in 3D and 2D cells. The ratio of all MUFA-PLs to all PUFA-PLs was calculated for each of the PC, PE, and PS lipid classes (**Figure S3G-I**). MUFA:PUFA ratios were significantly higher for PC, PE and PS lipids in 786-O and HT-1080 3D versus 2D cells. In NCI-H1299 cells, the MUFA:PUFA ratios were also higher in PC and PE lipids in 3D versus 2D cultures (**Figure S3G-I**). Taken together, these data showed distinct lipid profiles in 2D cultures, 3D cultures and tumors and were consistent with lipid changes as possible mediators of the observed differences in ferroptosis sensitivity.

### Increasing exogenous PUFAs re-sensitizes 3D cultures to ferroptosis

Peroxidation of PUFA-PLs is essential for ferroptosis induction^3,40,42^. Across all cell lines, PE-AA was less abundant in xenograft tumors and 3D cultures than 2D cultures (**Figure 2, S3**). Accordingly, we hypothesized that increasing AA levels would be sufficient to re-sensitize 3D cultures to ferroptosis. To test this hypothesis, we used cell lines with high 3D:2D RSL3 IC_50_ ratios (**Figure 1F**). Pretreatment of 3D HT-1080, NCI-H1299 and 786-O cultures with exogenous AA (10 μM, 16 h) re-sensitized these cells to RSL3 treatment (**Figure 3A, S4A**). 3D HEYA-8 and COR-L105 cultures were also sensitized to RSL3 treatment by AA pretreatment (**Figure S4A**). Moreover, the addition of exogenous AA to 3D cultures of 786-O sg*GPX4^iKO^* cells significantly enhanced cell death compared to DMSO-treated sg*GPX4* and sgCTL cells. Ferrostatin-1 (1 µM) cotreatment rescued cell death in AA pretreated 3D 786-O sg*GPX4^iKO^* cells, indicating that exogenous AA specifically enhanced ferroptosis sensitivity (**Figure 3B**). To test whether increasing endogenous AA was sufficient to re-sensitize 3D cultures to ferroptosis, we used the COX2 inhibitor DuP697. COX2 inhibition increases endogenous AA by inhibiting the conversion of AA to prostaglandin 2 (PGE2)^43^. Pretreatment with DuP697 (10 μM, 72 h) re-sensitized 3D HT-1080 and NCI-H1299 cultures to RSL3 (**Figure 3C**), albeit less so than exogenous AA (**Figure 3A**). Lower levels of PGE_2_ in the cell supernatant correlated with inhibition of AA conversion in DuP697-treated cells (**Figure S4B**).

**Figure 3.**
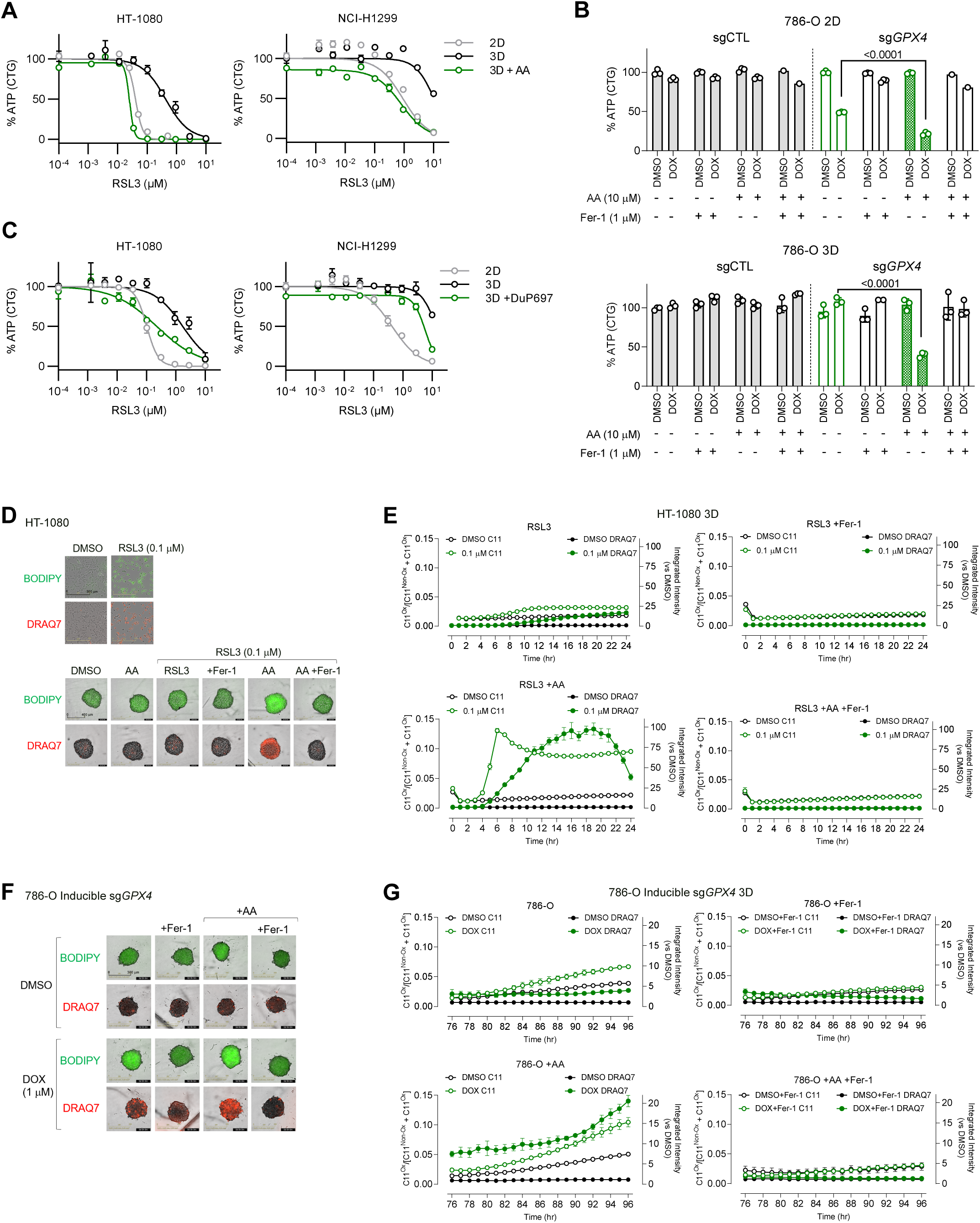
Increasing arachidonic acid sensitizes cells to ferroptosis induction in 3D cell culture. **A.** ATP levels (CTG) for 2D and 3D cultures of HT-1080 and NCI-H1299 cells, with AA (10 μM) added to the culture media at seeding, then treated with RSL3 (0-10 μM for 24 h). **B.** ATP levels (CTG) for 2D and 3D cultures of HT-1080 and NCI-H1299 cells pretreated with COX2 inhibitor (DuP697, 10 μM) at seeding, then treated with RSL3 (0-10 μM). **C.** ATP levels (CTG) for 2D and 3D cultures of 786-O inducible *GPX4* KO cells treated with doxycycline (1 µM/mL) for 48 h, then reseeded and treated with AA (10 μM). CTG measured 96 h post-doxycycline treatment. In **D** and **E,** images (**D**) and kinetics (**E**) of lipid peroxidation (BODIPY 581/591 C11) and cell death (DRAQ7) of HT-1080 3D cells pre-treated with AA (10 μM, 16 h) and ferrostatin-1 (1 μM), then RSL3. Images are of 6 h after RSL3 treatment. In **F** and **G**, lipid peroxidation (BODIPY 581/591 C11) and cell death (DRAQ7) of inducible 786-O 3D cells treated with doxycycline (1 µM/mL) for 48 h, then reseeded and treated with AA (10 μM). **D-G.** Kinetics of lipid peroxidation and cell death were quantified/imaged via IncuCyte. In **D** and **F**, image 10X magnification for 2D cells and 3D cells. In **E** and **G**, lipid peroxidation measured by C11^Ox^/(C11^NonOx^ + C11^Ox^) ratio at each time point. Cell death was determined in reference to DMSO (=1) at the same time point. Data represented three biological replicgates.

Cells undergoing ferroptosis accumulate lipid peroxides^44^. To determine whether the addition of AA to 3D cultures increased lipid peroxidation, we tracked the kinetics of lipid peroxidation and cell death using BODIPY 581/591 C11 and DRAQ7, respectively. BODIPY 581/591 C11 oxidation and DRAQ7 uptake were lower in 3D versus 2D HT-1080 cultures treated with RSL3 (**Figure 3D**). The addition of exogenous AA (10 μM, 16 h) to 3D HT-1080 cultures increased lipid peroxidation and cell death in response to RSL3 (**Figure 3D** and **E, bottom left**). BODIPY 581/591 C11 oxidation and DRAQ7 uptake were inhibited by Fer-1 (1 µM) co-treatment (**Figure 3E, top right** and **bottom right**). Lipid peroxidation was also increased in response to RSL3 in 3D NCI-H1299 and 3D HEYA8 cultures when AA (10 μM, 16 h) was added to the media (**Figure S4C**). Likewise, AA addition (10 μM, 16 h) increased cell death in 786-O sg*GPX4^iKO^* 3D cultures (**Figure 3F,G**). We further found that COX2 inhibition in 3D HT-1080 cultures accelerated cell death compared to vehicle-treated controls (**Figure S4D**). These results suggested that low PUFA levels may limit GPX4 sensitivity in 3D cultures.

### Enhanced MUFA metabolism promotes ferroptosis resistance in 3D conditions

MUFA-enriched membranes can protect cells from ferroptosis induction^5,23,24,27,40–42^. We wondered what mechanism explained the increased MUFA:PUFA ratio in 3D versus 2D cultures (**Figure S3G-I**). We hypothesized that differences in lipid metabolic gene expression between these conditions could underlie the shift in lipid content and ferroptosis resistance. Among lipid metabolism genes, we noted a significant upregulation of *SCD* expression in 3D versus 2D conditions for HT-1080, NCI-H1299 and 786-O cell lines (**Figure 4A**). SCD synthesizes the MUFAs palmitoleic acid (*cis*-9-C16:1) and OA (*cis*-9-C18:1) from palmitate (C16:0) and stearate (C18:0), respectively^39^. Consistent with the observed *SCD* upregulation, sterol regulatory element binding transcription factor 1 (*SREBF1*), the transcription factor that regulates *SCD* expression^45^, was also upregulated in these cells. Of note, *SCD* expression was not significantly increased in 3D versus 2D cultures for Calu-1 cells (**Figure 4A**). Correspondingly, these two cultures also showed no significant difference in C18:1 abundance in our lipidomic analysis (**Figure S5A**).

**Figure 4.**
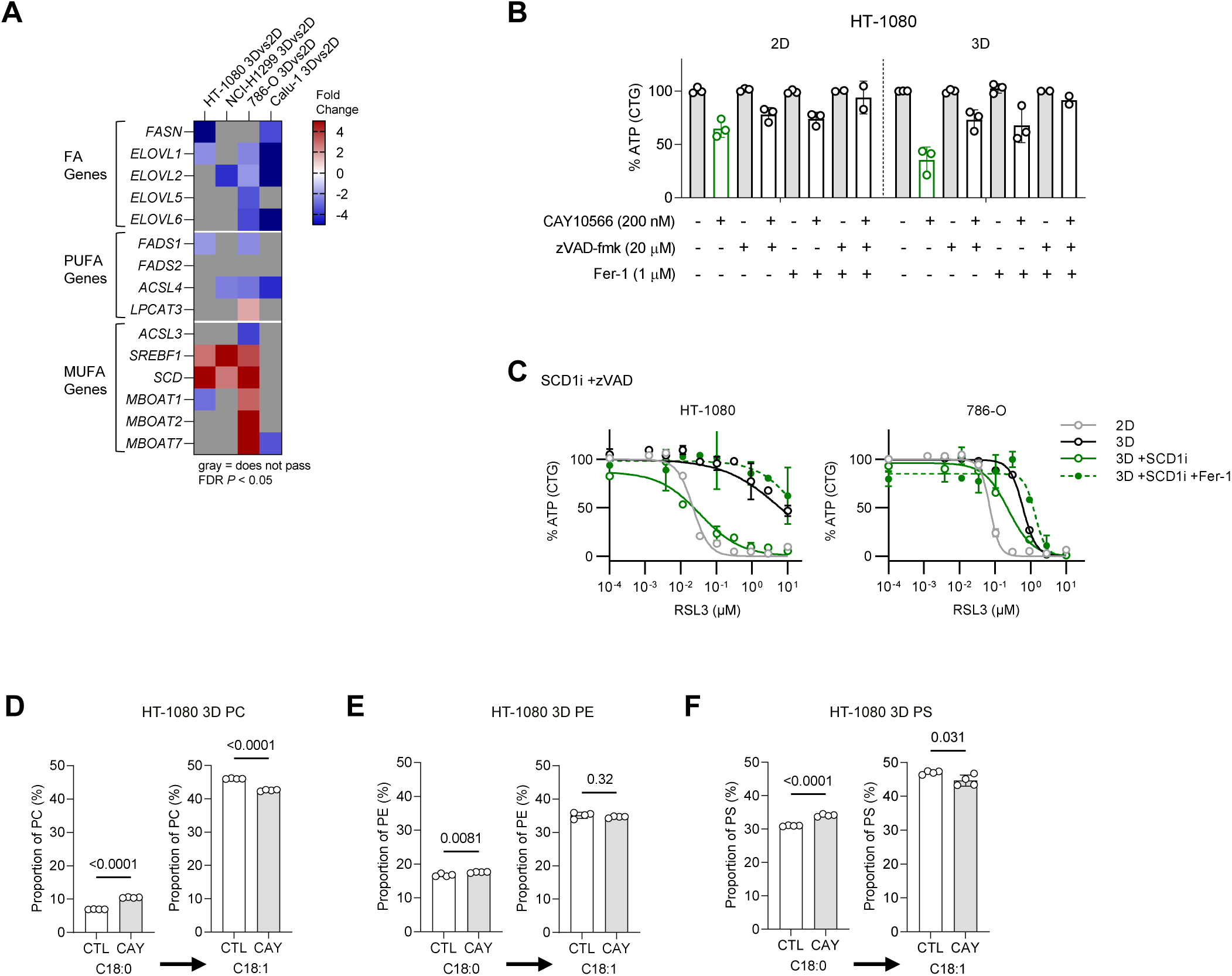
Decreasing MUFAs sensitize cells to ferroptosis induction in 3D cell culture. **A.** Baseline 3D vs 2D differentially expressed lipogenic genes (FDR *P* <0.05). Genome-wide expression measured via microarray (Clariom D assay). Gray indicates fold change was not significant (FDR *P* <0.05). Median of 4 biological replicates represented. **B.** ATP levels (CTG) for HT-1080 2D and 3D cultures treated with SCD1 inhibitor (CAY10566, 200 nM), zVAD (20 mM) and ferrostatin-1 (1 μM) added to cell media at seeding. Three biological replicates are shown. **C.** ATP levels (CTG) for 2D and 3D cultures treated with SCD1 inhibitor (CAY10566, 200 nM), zVAD (20 mM) and ferrostatin-1 (conc) added to cell media at seeding, then treated with RSL3. Biological replicates represented. **D**-**F.** Changes in 18:0-PL (SFA) ◊ 18:1-PL (OA) within PC (**D**), PE (**E**) and PS (**F**) in HT-1080 3D cells treated with *SCD1* inhibitor (CAY10566, 200 nM) for 16 hr. In **D**-**E**, data are presented as means ± SD and statistical significance was assessed using two-tailed unpaired Student’s t-test.

We hypothesized that SCD products, especially *cis*-9-C18:1, were necessary for ferroptosis resistance in 3D versus 2D cultures. We further hypothesized that decreasing MUFA content, thereby lowering the MUFA:PUFA ratio, would re-sensitize 3D cultures to ferroptosis induction. To test this hypothesis, we treated 3D and 2D cultures with the SCD inhibitor CAY10566 (SCDi, 200 nM). SCD is needed for survival in many cancer cell lines^46–48^ and SCD inhibition can induce apoptosis or ferroptosis^49,50^. Indeed, SCDi treatment caused HT-1080 cell death that was rescued by the pan-caspase inhibitor zVAD-fmk and Fer-1 in 3D and 2D conditions (**Figure 4B**). Accordingly, in 3D cultures, we used zVAD-fmk to isolate the effects of SCD inhibition on ferroptosis alone. SCDi treatment re-sensitized 3D HT-1080 and 786-O cultures to RSL3, with cell death under these conditions being prevented by Fer-1 (**Figure 4C**). SCDi treatment also significantly decreased the levels of C18:1-containing PLs in 3D HT-1080 cultures compared to vehicle-treated cells (**Figure 4D-F**). By contrast, SCDi treatment did not affect the proportion of the pro-ferroptotic PUFA-PLs PE-AA and PE-AdA in 3D cultures (**Figure S5B**). Thus, increased MUFA synthesis appears necessary for enhanced ferroptosis resistance in 3D versus 2D cultures.

### The anti-ferroptotic activity of MUFAs is structure-specific

Our results suggested that the MUFA:PUFA ratio dictated changes in ferroptosis sensitivity in 3D versus 2D cultures. Our SCD inhibitor studies showed that one or more products of SCD were found in greater abundance in 3D versus 2D cultures, thereby promoting ferroptosis resistance. SCD synthesizes both *cis*-9-C16:1 and *cis*-9-C18:1 (OA) which, together with endogenous fatty acid elongation pathways, can generate a diversity of final products^51^. Another enzyme, fatty acid desaturase 2 (FADS2), can also synthesize a distinct MUFA species (e.g., *cis*-6-C16:1, sapienate) that can be elongated to a *cis*-8-C18:1 product^52^ (**Fig. 5A**). Dietary uptake of both *cis* and *trans* MUFAs that are not synthesized endogenously can also alter the function of cancer and other cells^53,54^. Notably, different MUFA positional isomers cannot be distinguished by our mass spectrometry methods. Thus, it seemed valuable to further explore the MUFA structure-activity relationship with respect to ferroptosis inhibition.

**Figure 5.**
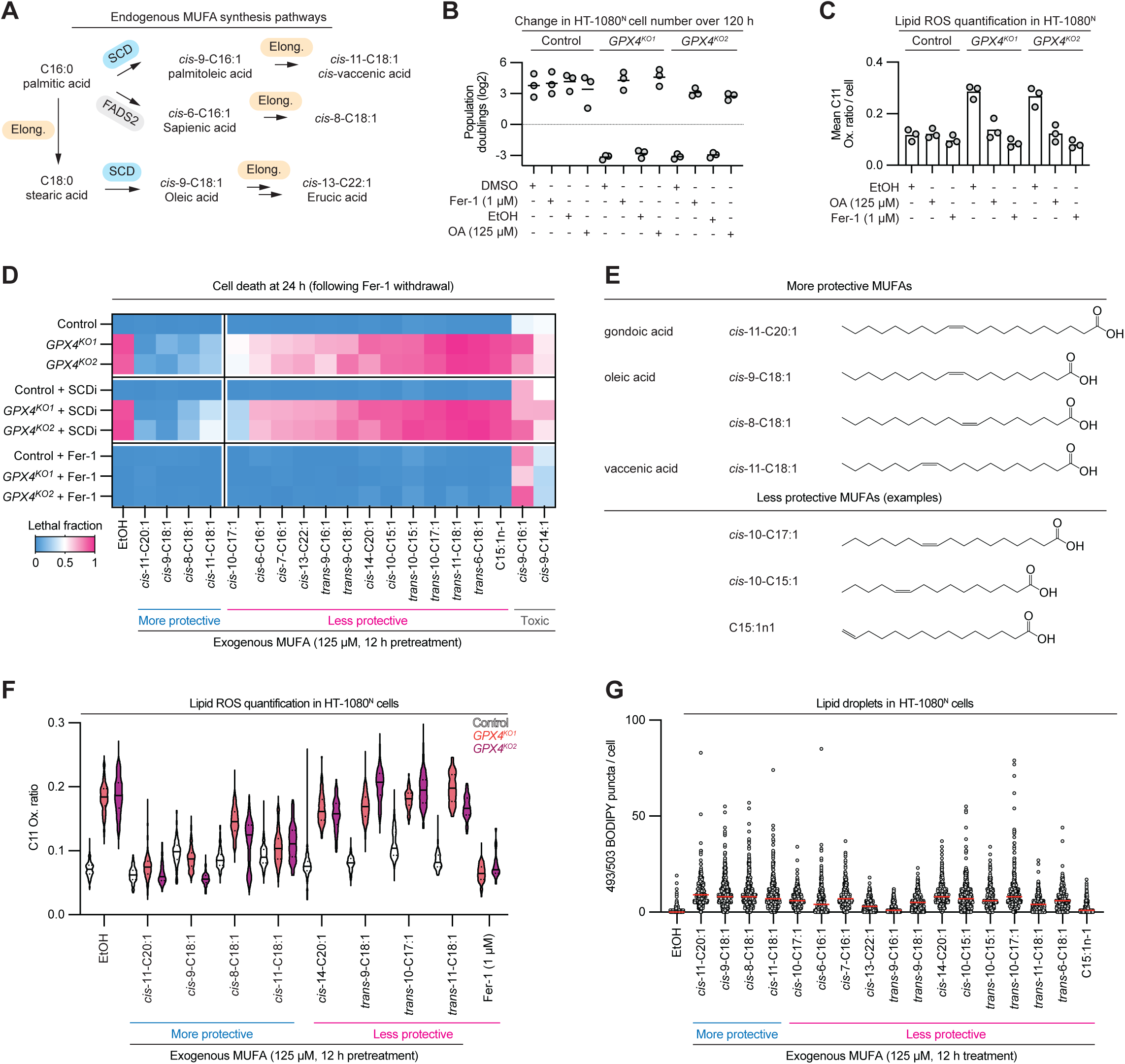
Ferroptosis inhibition is MUFA structure-specific. **A.** Outline of endogenous MUFA biosynthesis pathways. There are several endogenous C18:1 positional isomers and further elongation products. **B.** Cell death determined using imaging. *GPX4^KO1/2^* cells were grown in ferrostatin-1 (Fer-1, 1 µM) prior to the start of the experiment. Exogenous MUFAs were added 12 h before the removal of Fer-1 from *GPX4^KO1/2^* cultures. Heatmap values represent the mean of three independent experiments. **C.** Quantification of BODIPY 581/591 C11 (C11) fluorescence intensity (oxidized / non-oxidized) 2 h after Fer-1 withdrawal. Cells were pretreated with OA as in B. **D.** Cell death measured by integrating counts of live (mKate2-positive) and dead (SYTOX Green positive) cells. Cells were pretreated (12 h, 125 µM) with 19 different MUFAs prior to Fer-1 withdrawal. MUFAs were re-added at the time of Fer-1 withdrawal. **E**. Chemical structures of select protective and non-protective MUFAs. **F.** Quantification of C11 fluorescence intensity (oxidized / non-oxidized) 2 h after Fer-1 withdrawal (for *GPX4^KO1/2^* cultures). n = 43 – 256 individual cells quantified per condition across three independent experiments. Data indicate median and quartiles. **G**. Quantification of BODIPY 493/503 puncta per cell 12 h after MUFA treatment. Red horizontal bar is the median. n = 386 – 563 individual cells were counted per condition across three independent experiments.

For these experiments, we first employed two independent clonal HT-1080^N^ *GPX4* gene-disrupted (i.e., knockout, “KO”) cell lines that can only survive when ferrostatin-1 (Fer-1, 1 µM) is present in the culture medium^55^ (**Figure S6**). Note that the “N” superscript denotes cell lines engineered to express the live cell marker nuclear mKate2. Cells were also incubated with the dead cell marker SYTOX Green (SG). Counts of live (mKate2-positive) and dead (SG positive) cells were obtained by time-lapse imaging^56^. Consistent with previous results^5^, 2D HT-1080^N^ *GPX4^KO1/2^* cultures pre-incubated with exogenous *cis*-9-C18:1 (125 µM, 48 h) survived and proliferated upon Fer-1 withdrawal at similar rates to control cells (**Figure 5B**). C*is*-9-C18:1 pretreatment also reduced lipid peroxide accumulation in *GPX4^KO1/2^* cells upon Fer-1 withdrawal, as detected using BODIPY 581/591 C11 (**Figure 5C**). Thus, exogenous c*is*-9-C18:1 (OA) is sufficient to prevent ferroptosis in cells lacking *GPX4*.

We next tested the ability of 19 different endogenous, dietary, and other structurally distinct MUFAs (125 µM, 12 h pretreatment) to suppress ferroptosis in *GPX4^KO1/2^* cells upon Fer-1 withdrawal (summarized in **Figure 5D**). We observed structure-specific cell death protection. *Cis*-11-C20:1 (gondoic acid), *cis*-9-C18:1 (OA), *cis*-8-C18:1 and *cis*-11-C18:1 (*cis*-vaccenic acid) inhibited cell death by at least 50% in Fer-1-deprived *GPX4^KO1/2^* cells (**Figure 5E**). *Cis*-unsaturated MUFAs that were shorter and/or where the double bond was positioned further away from the middle of the molecule (e.g., *cis*-10-C17:1, *cis*-10-C15:1 and C15:1n-1) were less protective (**Figure 5D,E**). *Trans*-unsaturated MUFAs, including the OA isomer *trans*-9-C18:1 (elaidic acid), generally conferred little or no protection against ferroptosis (**Figure 5D**). The effects of *cis*-9-C16:1 (palmitoleic acid) and *cis*-5-C14:1 could not be evaluated in this experiment, as these MUFAs were toxic (**Figure 5D**). Incubation of MUFAs together with SCD1i (200 nM), to block the contribution of endogenous *cis*-9-C18:1, did not alter the observed patterns of cell death inhibition. Addition of Fer-1 (1 µM) rescued cell death in previously Fer-1-deprived *GPX4^KO1/2^* cultures, confirming that these cells were undergoing ferroptosis (**Figure 5D**).

We then further explored the mechanism of MUFA-mediated ferroptosis inhibition. We tested a subset of MUFAs for the ability to prevent BODIPY 581/591 C11 oxidation in HT-1080^N^ *GPX4^KO1/2^* cultures upon Fer-1 withdrawal. Consistent with the cell death results, incubation with the more protective MUFAs (as defined in **Fig. 5D**) *cis*-11-C20:1, *cis*-9-C18:1, *cis*-8-C18:1 and *cis*-11-C18:1 suppressed 581/591 BODIPY C11 oxidation while incubation with less protective MUFAs had less effect on probe oxidation (**Figure 5F)**. One possible reason why certain MUFAs could not inhibit ferroptosis was that they were not taken up into the cell and/or further metabolized. However, this did not appear to be a sufficient explanation for our results, as both more protective and less protective MUFAs stimulated neutral lipid accumulation and lipid droplet synthesis, as determined using BODIPY 493/503 staining, albeit to varying degrees (**Figure 5G**). Thus, diverse *cis*-unsaturated MUFAs are sufficient to inhibit ferroptosis in cells lacking GPX4 function.

### Protective MUFAs require ACSL3

Our lipidomic analyses of tumor xenografts, and 3D and 2D cell cultures, suggested that incorporation of MUFAs into PLs correlated with reduced ferroptosis sensitivity. Enzymes are required for MUFA incorporation into PLs. To protect from ferroptosis, exogenous OA requires enzymatic activation by acyl-CoA synthetase long-chain family member 3 (ACSL3), both in 2D culture and *in vivo*^5,23^ (**Figure 6A**). We hypothesized that *ACSL3* would be required for other MUFAs (125 µM, 12 h) to protect from RSL3-induced ferroptosis in HT-1080^N^ cells. To investigate, we employed established clonal Control and *ACSL3* gene-disrupted (“KO”) cell lines^5^, together with live/dead cell imaging. In addition to protective MUFAs identified previously (e.g., cis-9-C18:1, cis-8-C18:1), in this experiment we also observed that *cis*-13-C22:1, *cis*-6-C16:1 and *cis*-7-C16:1 achieved at least 50% cell death inhibition in Control cells (**Figure 6B**). *ACSL3* was generally required for the full protective effects of most MUFAs, albeit less so for *cis*-11-C20:1 (**Figure 6B**). Based on cell death phenotypes and BODIPY 493/503 neutral lipid staining (see below), it seems that *ACSL3^KO2^* cells harbor a stronger gene disruption than *ACSL3^KO1^* cells^5^. However, both clones exhibited similar behaviors (**Figure 6B**).

**Figure 6.**
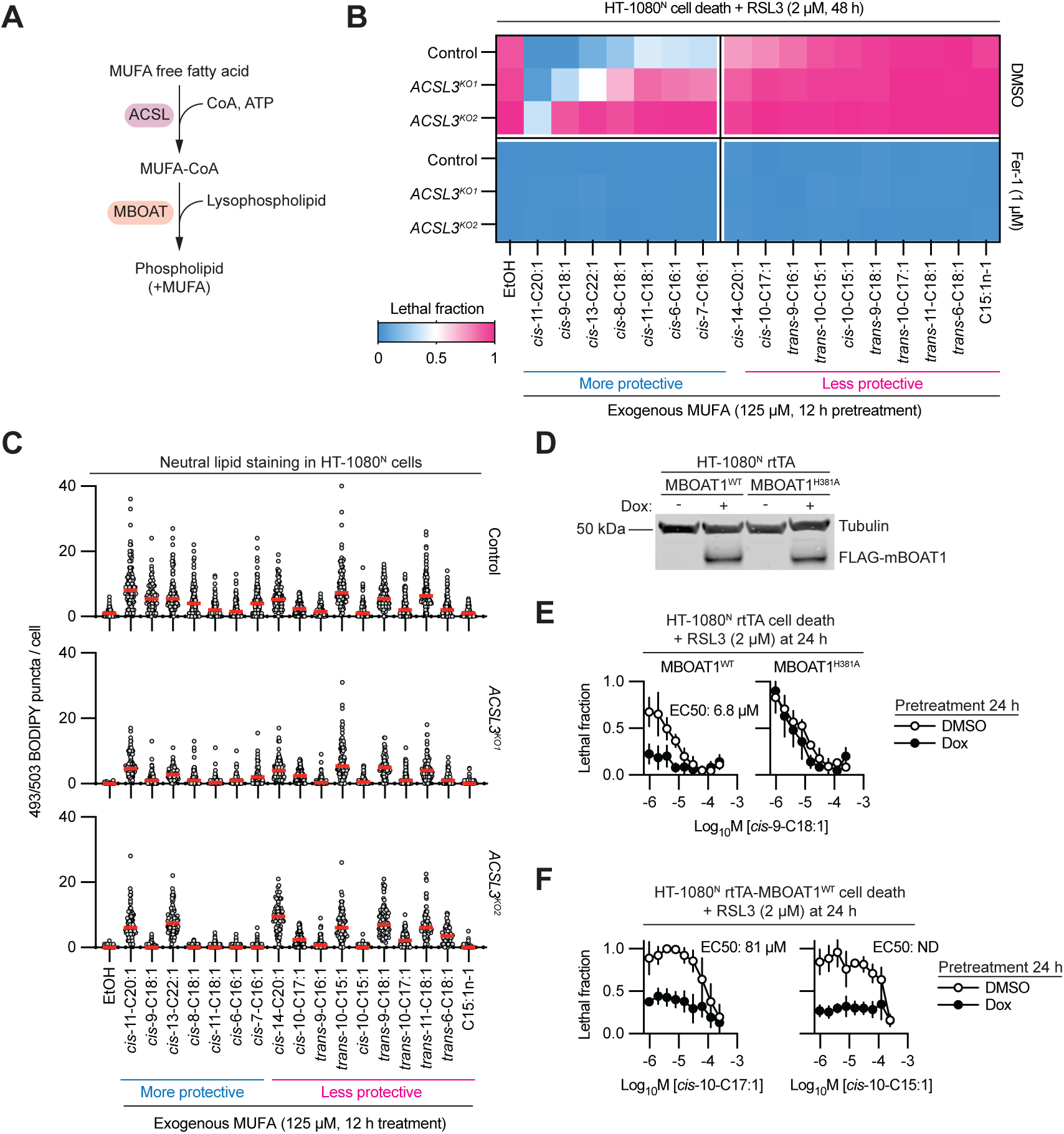
Ferroptosis inhibition is genetically regulated. **A.** Cartoon scheme of MUFA activation and incorporation into phospholipids. **B.** Cell death determined using imaging. *GPX4^KO1/2^*cells were grown in ferrostatin-1 (Fer-1, 1 µM) prior to the start of the experiment. Exogenous MUFAs were added 12 h before the removal of Fer-1 from *GPX4^KO1/2^*cultures. Heatmap values represent the mean of three independent experiments. **C.** Quantification of BODIPY 493/503 puncta per cell 12 h after MUFA treatment. Red horizontal bar is the median. n = 69 – 134 individual cells were counted per condition. **D.** Protein detection via immunoblotting. Doxycycline (Dox, 1 µg/mL, 24 h) was used to induce MBOAT1 expression. **E.** Cell death determined by imaging. **F.** Cell death determined by imaging. Data in E and F are mean ± SD from three independent experiments.

We next examined whether ACSL3-dependent protective effects correlated with the stimulation of neutral lipid accumulation. We hypothesized that MUFAs which exhibited *ACSL3*-dependent protective effects would also exhibit *ACSL3*-dependent stimulation of neutral lipid synthesis. This hypothesis was supported, in general (**Figure 6C**). Thus, *cis*-9-C18:1, *cis*-8-C18:1, *cis*-11-C18:1, *cis*-6-C16:1 and *cis*-7-C16:1 had less ability to stimulate neutral lipid accumulation in *ACSL3*-mutant cells, coincident with the loss of anti-ferroptotic activity (**Figure 6B**, **C**). By contrast, *cis*-11-C20:1 (gondoic acid), retained the ability to stimulate substantial neutral lipid accumulation in *ACSL3^KO1/2^*, coincident with the ability to suppress ferroptosis (**Figure 6A**, **B**). We infer that *cis*-11-C20:1 can be activated in part by a different ACSL enzyme. Some less protective MUFAs (e.g., *cis*-10-C15:1) had a reduced ability to stimulate neutral lipid synthesis in *ACSL3^KO1/2^* cells versus control cells, while other less protective MUFAs (e.g., *cis*-14-C20:1) could stimulate lipid droplet accumulation in an *ACSL3*-independent manner (**Figure 6B**, **C**). Thus, *ACSL3*-dependent protection against ferroptosis did not correlate strictly with the *ACSL3*-dependent stimulation of neutral lipid synthesis.

Our results suggested that certain less protective MUFAs could be activated (i.e., conjugated to CoA, a necessary step in further metabolism), yet were still unable to suppress ferroptosis. We hypothesized that this failure to suppress ferroptosis may be due to decreased incorporation into membrane PLs. To investigate, we made use of HT-1080^N^ cells engineered to overexpress wild-type or predicted catalytically inactive (H381A) lysophospholipid acyltransferase MBOAT1 in response to Dox treatment^21^ (**Figure 6D**). As expected, the more protective MUFAs *cis*-9-C18:1, *cis*-11-C18:1 and *cis*-8-C18:1 were potent inhibitors of RSL3-induced ferroptosis, even in the absence of Dox treatment (EC_50_s = 6.8 µM, 16 µM and 21 µM, respectively) (**Figure 6E**, **S7A**). Expression of the MBOAT1^H381A^ mutant had no effect on the ability of *cis*-9-C18:1 to inhibit ferroptosis (**Figure 6E**). The less protective MUFAs *cis*-10-C17:1 (EC_50_ = 81 µM) and *cis*-10-C15:1 (EC_50_ could not be determined) were, as expected, less potent suppressors of RSL3-induced ferroptosis in vehicle-treated MBOAT1^WT^ cells. However, the protective effects of Dox-induced MBOAT1^WT^ overexpression could be further enhanced by high concentrations of *cis*-10-C17:1 or *cis*-10-C15:1 (**Figure 6F**). Thus, the ability of certain MUFAs to protect against ferroptosis may be limited by MBOAT1 activity.

### MUFAs as probes for ferroptosis-essential PUFAs

Our cell death and lipidomic analyses suggested that displacement of more oxidizable PUFAs (e.g., AA) by less oxidizable MUFAs (e.g., C18:1) correlated with and likely was sufficient to drive ferroptosis inhibition in xenograft tumor and 3D growth conditions versus 2D growth conditions. We recognized an opportunity to use our MUFA analogs as mechanistic probes to globally define lipid species that were selectively perturbed by “more protective” versus “less protective” MUFAs. We incubated HT-1080^N^ cells grown in 2D culture with the protective MUFAs *cis*-9-C18:1, *cis*-11-C18:1 and *cis*-8-C18:1 and the less protective MUFAs *cis*-10-C17:1, *cis*-10-C15:1 and C15:1n-1 (all at 125 µM, 12 h), then performed shotgun lipidomic analysis (**Figure 7A**).

**Figure 7.**
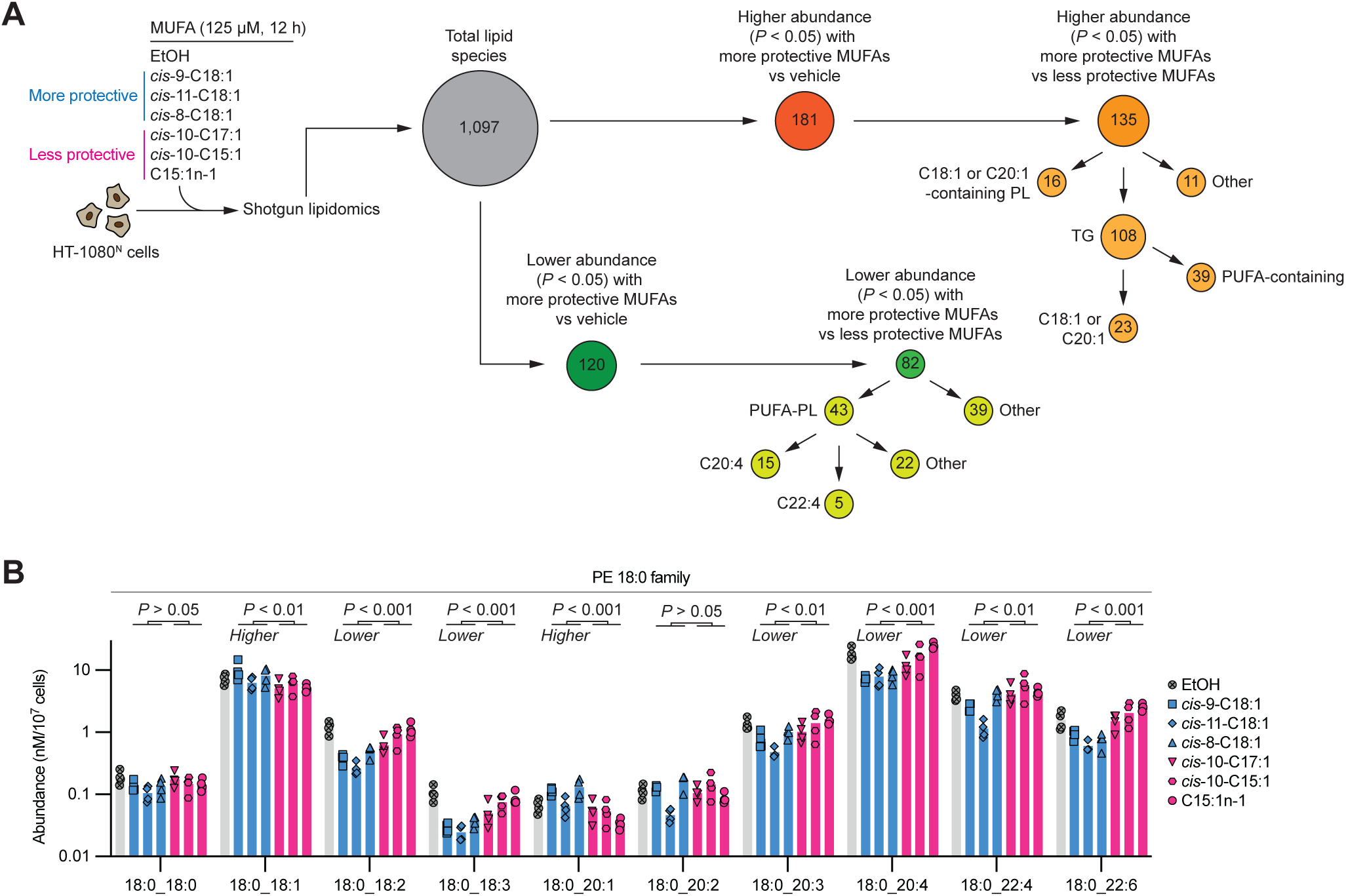
Exogenous MUFAs as probes for ferroptosis regulators. **A.** Outline of the shotgun lipidomic analysis and breakdown of altered lipid species. **B.** Lipid abundance for the PE 18:0 family of lipid species. Individual datapoints from four separate experiments are shown.

We identified 1,097 lipid species in at least one condition. 120 lipid species had significantly (*P* < 0.05) reduced abundance compared to the vehicle (ethanol) control in response to all three protective MUFAs, of which 82 also were found at significantly lower abundance in cells treated with more protective MUFAs than with less protective MUFAs. Of these 82 species, 52% (43/82) were PUFA-PLs, including 15 species that contained C20:4 (AA) and five species that contained C22:4 (AdA). By contrast, 181 lipids were significantly increased in cells incubated in more protective MUFAs versus vehicle control, of which 135 were significantly altered by more protective MUFAs than by less protective MUFAs. Of these lipids, 80% (108/135) were TGs, 23 of which were enriched with C18:1 species and 39 of which contained at least one PUFA chain (**Figure S7B**). Thus, protective MUFAs both reduced the abundance of oxidizable PUFA-PLs and increased the abundance of PUFA-loaded TGs.

To further illustrate the relative impacts of more protective versus less protective MUFAs on the lipidome we focused on the family of PE lipids containing C18:0 at one position (**Figure 7B**). We obtained reliable data for 10 species in this class. Of these ten lipids, eight were significantly altered in abundance between more protective and less protective MUFAs; all six PUFA-PEs, including C18:0_20:4 and C18:0_22:4, had significantly lower average abundance following treatment with more protective than less protective MUFAs (*P* <0.05). These results were consistent with the changes in lipid composition observed in 3D versus 2D culture conditions, suggesting that the loss of PUFA-PEs may underlie ferroptosis resistance in a broad range of conditions.

## DISCUSSION

GPX4 has been proposed as a candidate target for ferroptosis therapies. Early studies indicating that direct injection of RSL3 at the site of tumor cell implantation could prevent tumor formation^10^. More recently, new small molecule GPX4 inhibitors with superior pharmacokinetic properties have been developed, with mixed results following systemic administration *in vivo*. One inhibitor, compound 24, was tolerated in mice and showed partial target engagement *in vivo* but had no effect on diffuse large B cell lymphoma xenograft tumor growth (notably, despite strong effects on these same cells in 2D culture)^22^. An orally bioavailable GPX4 inhibitor, compound 28, engaged GPX4 strongly in mouse HT-1080 xenograft tumors, but only slowed tumor growth significantly in combination with an inhibitor of cyclin dependent kinases 4 and 6, and caused noticeable body weight loss^21^. Thus, the therapeutic window for single agent GPX4 inhibitor strategies remains unclear and may be limited by on-target toxicities. Our findings suggest that lipid-mediated ferroptosis resistance mechanisms activated in non-adherent cells could act to further limit the size of this window.

We find that xenograft tumors *in vivo*, and 3D cell cultures *in vitro*, can be more resistant to genetic and chemical inhibition of GPX4 than 2D cultures of the same cells. This resistance is not explained by higher expression of *GPX4* itself, *FSP1*, or other enzymes that confer ferroptosis resistance by synthesizing radical trapping antioxidants. Rather, increased expression of the lipid metabolic gene *SCD* appeared to drive a MUFA-centric metabolic program that conferred protection against ferroptosis, at least in 3D versus 2D cultures. We propose that a higher MUFA:PUFA ratio limits the oxidizability of the membrane, preventing the accumulation of membrane lipid peroxides to lethal levels. High cell confluence has been suggested to reduce ferroptosis sensitivity in epithelial cells, in part, through an E-cadherin (*CDH1*)-based signaling mechanism^57^. Whether activation of the E-cadherin pathway can also alter lipidomic profiles is currently unknown.

3D models such as spheroids and organoids are considered an intermediate system between 2D cell culture and *in vivo* tumors^35^. Characterization of this intermediate system has shown that cells in 3D culture often experience more oxidative stress than cells in 2D culture^58,59^. However, we find that *SCD* upregulation and greater MUFA abundance can overcome this potential increase in oxidative stress, at least in response to lipid peroxidation. The specific features of the 3D environment that lead to *SCD* upregulation will require further investigation. Other studies have found differences in metabolism and ferroptosis sensitivity between cells residing at different positions within the spheroid (i.e., core versus the outer layer)^58^. Future studies will also be needed to elucidate whether such differences are relevant in our cancer cell models.

Our SCD data suggest an important role for SCD products like *cis*-9-C18:1 in ferroptosis resistance in 3D environments. However, cancer cells can synthesize or acquire from the environment a variety of structurally distinct C18:1 and other MUFA species. Importantly, many common mass spectrometry methods cannot distinguish between different C18:1 positional isomers. In general, we find that *cis*-unsaturated MUFAs were ferroptosis-suppressive while *trans*-unsaturated MUFAs were not, despite evidence of equivalent uptake into the cell. The differential ability of *cis* versus *trans* MUFAs to inhibit ferroptosis may be linked to their effects on membrane structure; lipids with the kinked *cis* bond could have distinct interactions with PL bilayers compared to lipids with a straight *trans* bond. However, such differences seem unlikely to explain our results, as fully saturated fatty acids can also inhibit ferroptosis in some models^25,60^. Rather, we propose that *cis*-MUFAs are preferred substrates for downstream MBOAT-family lysophospholipid acyltransferases, which can use these substrates to reacylate lyso-PLs preferentially with MUFA species in competition with or at the direct expense of PUFA species^61^. Indeed, only protective *cis*-MUFAs reduced the abundance of various ferroptosis-relevant PE-PUFA species.

The experimental results described here have certain limitations. For sg*GPX4^iKO^* tumor xenografts, less than 100% GPX4 protein elimination could explain the lack of effect on tumor growth. It is possible that reducing tumor growth via *GPX4* disruption requires complete elimination of this protein. This data also indicates that even with significant reduction of GPX4 protein, it is not required for growth in established tumors. Additionally, our survey of the effects of different MUFAs on ferroptosis is not exhaustive; it is possible that some MUFA species not examined here can inhibit ferroptosis and/or do not act through the proposed *ACSL3*-dependent mechanism. Indeed, *cis*-11-C20:1 (gondoic acid) was able to suppress ferroptosis even in the absence of *ACSL3* and may therefore act through a distinct lipid metabolic pathway. More generally, our findings indicate that generalizing GPX4 inhibitor sensitivities obtained in 2D models to more complex 3D environments or *in vivo* needs to be performed with caution. Our results may, however, provide a rationale for therapeutic strategies that combine GPX4 inhibitors with SCD inhibitors to restore “2D-like” ferroptosis sensitivity *in vivo*.

## ACKNOWLEDGEMENTS

We thank AbbVie’s Genomics Research Center, G. Forcina, L. Magtanong, S. Petiwala and A. Neely for experimental assistance, K. Fujihara and N. Clemons for advice, and L. Murray, W. Lee, S. Manukian, A. Gautam, L. Leak, J. Olzmann, M. Jouni, E. Nyamugenda and D. Verduzco for comments on this work. This work was supported by the NIH (GM122923 to S.J.D.) and funds from AbbVie, Inc.

## AUTHOR CONTRIBUTIONS

Conceptualization, V.S.P., L.E.P., J.I., J.P., S.J.D., D.C.P, R.P.; Methodology, V.S.P., L.E.P., J.I., G.A.A., J.P., E.Y.A., A.C., P.N.; Investigation, V.S.P., L.E.P., J.I., G.A.A., J.P., E.Y.A., S.J., S.S., A.C., P.N.; Writing - Original Draft, V.S.P., L.E.P., D.C.P., S.J.D., R.P.; Resources, S.J.D., D.C.P., R.P.; Supervision, S.K., D.C.P, S.J.D., R.P.

## ABBVIE DISCLOSURE STATEMENT & DECLARATION OF INTERESTS

V.S.P., J.P., S.J., S.S., A.C., P.N., S.K., D.C.P. and R.P. are employees of AbbVie and may own stock. J.I. was an employee of AbbVie at the time of the study. S.J.D., L.E.P., G.A.A. and E.Y.A are affiliated with Stanford University. The design, study conduct, and financial support for this research were provided by AbbVie. AbbVie participated in the interpretation of data, review, and approval of the publication. S.J.D. is an inventor on patents related to ferroptosis.

## MATERIALS & METHODS

### Statement on animal care/guidelines

All protocols were approved by the Institutional Animal Care and Uses Committee (IACUC) at AbbVie Inc. and performed in accordance with National Institutes of Health Guide for the Care and Use of Laboratory Animals. All animals were housed under standard laboratory conditions with *ad libitum* access to food and water in a temperature and humidity-controlled room with a 12:12 h light:dark cycle. Female SCID and SCID-Beige mice with starting age ranging from 6-8 weeks (20 g, Charles River Laboratories, Wilmington, MA, USA) were housed in groups of ten in solid bottom Plexiglas cages equipped with constant air circulation in an Association for Assessment and Accreditation of Laboratory Animal Care International (AAALAC) approved facility. Animals were acclimated to the laboratory environment for 5-7 d before the study.

### CDX tumor mouse experiments

Commercially available doxycycline diet (Inotiv TD.01306, Madison, WI) containing a dose of 625 mg/kg was continuously fed to the mice starting 7 d prior to the initiation of the study and continued for the duration of the study. Mice were inoculated subcutaneously with 5 x 10^6^ parental, NTC or GPX4 786-O (SCID Female), Calu-1 (SCID-Beige Female) and NCI-H23 (SCID-Beige Female) cells at 0.1 mL/mouse in a mixture of 1:1 cell medium and Matrigel (BD Biosciences, Bedford, MA). Tumor volumes were determined using a caliper ellipsoid model: L x W^2^/2, where the larger (L) and smaller (W) perpendicular dimensions are measured.

### DepMap analysis

Analyses were performed using the DepMap (23Q4) dataset accessed in January 2024 (https://depmap.org/portal/download/). Data points were downloaded and plotted in Prism.

### Cell lines and culture conditions

786-O (CRL-1932, gender: male), Calu-1 (HTB-54, gender: male), NCI-H23 (CRL-5800, gender: male), NCI-H1299 (CRL-5803, gender: male), A549 (CCL-185, gender: male), HT-1080 (CCL-121, gender: male), 293T (CRL-3216), PANC1 (CRL-1469 gender: male) and T98G (CRL-1690 gender: male) cell lines were obtained from ATCC, expanded, frozen into aliquots and used for subsequent experiments. COR-L105 (92031918, gender: male) cell line was obtained from ECACC. HT-1080^N^ Control and *ACSL3^KO1/2^* cells were previously described^5^. HT-1080^N^ Control and *GPX4^KO1/2^* cells were described^55^. HT-1080^N^ rtTA-MBOAT1^WT^ and MBOAT1^H381A^ cells were described^21^. 786-O, Calu-1, NCI-H23, CORL-105, HEYA-8 and NCI-H1299 cells were cultured in RPMI 1640 medium (Cat# 11875093, Gibco) and supplemented with 10% fetal bovine serum (FBS, Cat# 16140071, Gibco), 1% sodium pyruvate (Cat# 11360070, Gibco) and 1% Pen/Strep (details). Medium supplemented with 10% tetracycline-approved FBS (Cat# 631106, Takara) was used with Dox-inducible cell lines. For other experiments, HT-1080, A549, 293T, PANC1 and T98G cells were cultured in Dulbecco’s modified Eagle medium (DMEM, Cat# 11995065, Thermo Fisher Scientific, Waltham, MA) supplemented with 10% fetal bovine serum (FBS, Cat# 26140-079, Gibco), and 0.5 U/mL Pen/Strep (P/S, Cat# 15070-063, Gibco). HT-1080 cell lines were additionally supplemented with 1% MEM nonessential amino acids (Cat# 11140050, Life Technologies). 1X PBS (Cat# 97062-338, VWR) and trypsin (Cat# 25200114, Gibco) were used for standard tissue culture experiments. For these cell lines, cells were counted using a Cellometer Auto T4 cell counter (Nexcelom, Lawrence, MA, USA).

### Chemicals and reagents

Erastin2 and ML162 were synthesized by Acme (Palo Alto, CA). RSL3 was obtained from Selleck Chemical (Cat# S8155). Doxycycline (Sigma, Cat# D3072), ferrostatin-1 (Cat# SML0583) and dimethyl sulfoxide (DMSO) (Cat# 276855) were from Sigma-Aldrich. BODIPY 581/591 C11 (Cat# D3861) and BODIPY 493/503 (Cat# D3922) were from ThermoFisher. The following fatty acids were obtained from different suppliers: arachidonic acid (Sigma, Cat# 10931), cis-6-hexadecenoic acid (*cis*-6-C16:1, Cayman Chemical, Cat# 9001845), cis-7-hexadecenoic acid (*cis*-7-C16:1, Cayman Chemical, Cat# 10007290), cis-8-octadecenoic acid (*cis*-8-C18:1, Cayman Chemical, Cat# 27447), cis-10-heptadecenoic acid (*cis*-10-C17:1, Cayman Chemical, Cat# 19748), trans-10-heptadecenoic acid (*trans*-10-C17:1, Santa Cruz Biotechnology, Cat# sc-507081), 10(Z)-pentadecenoic acid (*cis*-10-C15:1, Cayman Chemical, Cat# 22362), 10(E)-pentadecenoic acid (*trans*-10-C15:1, Cayman Chemical, Cat# 22467), cis-11-eicosenoic acid/11(Z) eicosenoic acid (*cis*-11-C20:1, Sigma-Aldrich, Cat# 44878 or Cayman Chemical, Cat# 20606), 13(Z)-docosenoic acid (*cis*-13-C22:1, Cayman Chemical, Cat# 90175 or Sigma-Aldrich, Cat# E3385), 14-pentadecenoic acid (C15:1n-1, Cayman Chemical, Cat# 19724 or Sigma-Aldrich, Cat# AMBH93E4C3FF), 14(Z)-eicosenoic acid (*cis*-14-C20:1, Cayman Chemical, Cat# 10009375), cis-vaccenic acid (*cis*-11-C18:1, Cayman Chemical, Cat# 20023), trans-petroselinic acid (*trans*-6-C18:1. GLPBio, Cat# GC18802), myristoleic acid (*cis*-9-C14:1, Sigma-Aldrich, Cat# M3525), palmitelaidic acid (*trans*-9-C16:1, TargetMol, Cat# T19500), palmitoleic acid (*cis*-9-C16:1, Cayman Chemical, Cat# 10009871), oleic acid (*cis*-9-C18:1, Caymen Chemical, Cat# 90260), elaidic acid (*trans*-9-C18:1, Sigma-Aldrich, Cat# E4637) and trans-vaccenic acid (*trans*-11-C18:1, Cayman Chemical, Cat# 15301).

### Generation of inducible GPX4 gene-disrupted cell lines

786-O, Calu-1 and NCI-H23 Cas9 stably expressed cell lines were plated in a 6-well plate at 2.0 x 10^5^ cells seeding density. The next day, varying volumes of lentivirus (25 mL, 10 mL, puro alone, and untreated) and 2 mL of polybrene were added to each well. After 24 h, medium containing virus was removed and fresh medium with puromycin (5 mg/mL) was added. After 48 h of puromycin treatment, the well with 35-50% viability was expanded and maintained in puromycin (1 µg/mL). sgRNA for *GPX4*: AGAGATCAAAGAGTTCGCCG and Control: GGCAGTCGTTCGGTTGATAT.

### Genome-wide 3D CRISPR screen

The Humagne library sets (C & D) used for the screen were designed for enAsCas12 as previously described^62^. The plasmid libraries were transfected into HEK 239T cells to produce lentiviral pools and subsequently transduced into the NCI-H1299 enCas12a cell line^34^. Cells were infected with libraries at a multiplicity of infection of 0.3 and selected with puromycin (3 μg/ml) for 3 d. Aliquots were saved as day 0 stocks and remaining cells were seeded into 2D monolayer cultures or 3D spheroids in Corning Elplasia 12K flasks (∼100 cells/microcavity). The screen was performed at ∼1,000X cell number coverage per sgRNA. For the 3D RSL3 treatment arm, cells were treated with RSL3 (2 µM) throughout the screen for 9 d. Each arm was performed with 4 biological replicates. Genomic DNA was extracted with Machery-Nagel NucleoSpin Blood XL (Takara, Cat#740950). sgRNAs were PCR amplified from gDNA (Clontech, Takara, Cat#639242) and measured with deep sequencing on NovaSeq (Illumina).

Depletions and enrichments of sgRNA between end time point of 3D DMSO and 3D RSL3-treated samples were used to calculate gene effect scores. For analysis, sequencing data from each CRISPR screen were demultiplexed and reads with sgRNA sequences were quantified using custom Perl scripts^34^. Sequences in between the flanking regions of gRNAs were extracted and then mapped to the Humagne set (C & D) library. Only sequences with no mismatches were used in the calculation of guide-level read counts. Through the edge R Bioconductor package, reads were normalized between samples using the trimmed mean of M values (TMM) method. To measure guide heterogeneity across timepoints the Gini index for each sample was calculated. Differential analysis of guide level counts was performed using the limma R Bioconductor package. All guides per gene are on the same construct and so the guide-level log fold change is synonymous with gene level effect. Statistical tests of the differential analysis were carried out using the Robust Rank Aggregation (RRA) method at the gene level.

### Western blotting

For inducible *GPX4* KO cell lines, cells were lysed RIPA buffer containing phosphatase/protease inhibitor (HALT Protease and Phosphatase Inhibitor Cocktail, Thermo, Cat# 78438) and EDTA on ice and spun down at 13000 rpm for 10 m. Protein concentration of supernatant was measured using Pierce BCA Protein Assay (Thermo, Cat# 23225) as per protocol, then boiled at 95°C for 5 m. Protein (50 mg) was run on a 4-12% gradient NuPAGE Bis-Tris Gel (Invitrogen, Cat# NP0321) and transferred to PVDF blots on iBlot 2 (Invitrogen). Blots were incubated with blocking buffer for 30 m, washed and incubated with primary antibody (1:2000 dilution) and actin (1:5000) at 4°C overnight. After 3 wash steps, secondary antibody was incubated for 1-2 h at RT. Blots were imaged with Odyssey.

For some experiments, 500,000 cells/well were seeded into 6-well plates (Thermo Fisher Scientific, Cat# 07-200-83) in 2 mL medium. After 4 h, cells were washed once with 1X PBS (VWR, Cat# 97062-338), then scraped into a small volume of PBS. Cells were pelleted by spinning cells at 900 x *g* for 2 min. Pellets were lysed with 50 µL of 9 M urea (Sigma Aldrich, Cat# U5378). Lysates were subsequently sonicated and spun for 15 min in a centrifuge at max speed. Supernatants were collected and transferred to a fresh 1.5 mL Eppendorf tube, and protein abundance was quantified with the BCA assay. 30 µg of protein from each lysate was loaded onto a Bolt 4-12% Bis-Tris Plus SDS gel (Life Technologies, Cat# NW04120BOX) for separation for 75 min at 100 V. The gel was transferred to a nitrocellulose membrane using the iBlot2 system (Life Technologies). Membranes were probed with primary antibody overnight in a hybridization bag at 4°C. Antibody designations and dilutions (and suppliers) are as follows: anti-α-tubulin DM1A 1:5000 (Thermo Fisher Scientific, Cat# MS581P1); anti-GPX4 EPNCIR144 1:1000 (Abcam, Cat# ab125066). Donkey anti-rabbit secondary antibody (IRDye 800CW, LI-COR Biosciences, Cat# 926-32213) and donkey anti-mouse antibody (IRDye 680LT, LI-COR Biosciences, Cat#926-68022) were used at 1:15,000 dilution to visualize bands. Membranes were imaged using the LI-COR CLx Imaging System.

### Cell viability measurements

For 2D cell culture viability measurements, 2,000 cells were seeded in 384-well plates (Corning, Cat# CLS3825). After 24 h cells were treated with RSL3 (0-10 μM). ATP production was measured with CellTiter-Glo assay (Promega, Cat#G7570) as per manufacturer’s instructions. For 3D cell culture, spheroids were allowed to form for 48 h before RSL3 treatment. ATP production was measured with CellTiter-Glo 3D assay (Promega, Cat#G9681) as per manufacturer’s instructions. For the inducible cell lines, doxycycline was added in 2D cell culture, then after 36 h post-dox, seeded as 3D spheroids in ULA plates. CTG was performed in 2D (96 h post dox) and 3D (120 h post dox).

### Cell death experiments

For HT-1080^N^ Control and *GPX4^KO1/2^* cell death measurement, ferrostatin-1 was washed out of plates 24 h after seeding and medium was replaced with SYTOX Green (20 nM)-containing medium with or without ferrostatin-1 (1 μM). After 8 h, cells were imaged in phase contrast and green channels. Each well was imaged as described below, then automated object detection was performed in parallel to data acquisition using the Zoom software package (V2016A/B) using a routine with the following settings to count SG^+^ objects (parameter adaptive, threshold adjustment: 3; edge split on; edge sensitivity 0). Cell death was quantified as number of SG^+^ objects per mm^2^.

For analysis of MUFA effects in HT-1080^N^ Control and *GPX4^KO^*^1/2^ cells, 2,500 HT-1080^N^ Control, *GPX4^KO1^*, or *GPX4^KO2^*cells were seeded into 200 μL a 96 well plate (Corning, Cat# 3610) into 200 μL media containing ferrostatin-1 (1 μM) and briefly centrifuged to settle cells. 24 h after seeding, medium was removed from plate and media with fatty acid at 125 μM as well as ferrostatin-1 (1 μM) were added to wells. 12 h after fatty acid treatment, media was removed from plate and cells were washed gently with 200 μL PBS. PBS was removed and replaced with 200 μL media containing 20 nM Sytox green and either DMSO, erastin2 (1 μM), or ferrostatin-1 (1 μM). For experiments in *ACSL3^KO1/2^* cell lines, on day 0, 750 HT-1080^N^ Control or *ACSL3^KO1/2^* cells were seeded into a 384 well plate. 24 h later, cells were pretreated with 125 μM fatty acid in a total volume of 50 μL. 12 h later, media was removed and media containing SYTOX Green (20 nM) with vehicle control (DMSO) or RSL3 (2 μM), and with vehicle control (DMSO) or ferrostatin-1 (1 μM), was added to the plate.

Cell death was analyzed using the scalable time-lapse analysis of cell death kinetics (STACK) technique. After 12 h, 24 h, and 48 h of incubation, mKate2^+^ (live cells) and SG^+^ (dead cells) objects were counted using the Essen IncuCyte Zoom (Essen BioScience, Ann Arbor, MI). For all cell death experiments, each well was imaged (1392 x 1040 pixels at 1.22 μm/pixel) using a 10x objective lens in phase contrast, green fluorescence (ex: 460 +/- 20, em: 524 +/- 20, acquisition time: 400 ms) and red fluorescence ((ex: 585 ± 20, em: 665 ± 40, acquisition time: 800 ms). Automated object detection was performed in parallel to data acquisition using the Zoom software package (V2016A/B). The following settings were used for HT-1080^N^ cells to count mKate2^+^ objects: parameter adaptive, threshold adjustment 0.5; edge split on; edge sensitivity -28; filter area min 100 µm^2^; filter area max 600 µm2; eccentricity max 0.9), SG^+^ objects: parameter adaptive, threshold adjustment: 5; edge split on; edge sensitivity 0; and overlap (e.g. mKate2^+^ and SG^+^) objects: filter area, min 90 μm^2^. For PANC1^N^ and T98G^N^ cells the following setting were used for mKate2^+^ objects: adaptive threshold adjustment 0.5; edge split on; edge sensitivity -15; filter area min 76 μm^2^, maximum 600 μm^2^; eccentricity max 0.9, for SG^+^ objects: adaptive threshold adjustment 5.0; edge split on; edge sensitivity 0, and for overlap objects: filter area min 90 μm^2^. For experiments in *ACSL3^KO1/2^* cell lines, the following detection setting were used. For mKate2^+^ objects: parameter adaptive, threshold adjustment 0.5; edge split on; edge sensitivity -28; filter area min 100 μm^2^; filter area max 600 μm^2^; eccentricity max 0.9, SG^+^ objects: parameter adaptive, threshold adjustment: 5; edge split on; edge sensitivity 0), and overlap objects (e.g. mKate2^+^ and SG^+^) objects (filter area min 90 μm^2^). Counts of live (mKate2^+^), dead (SG+) and overlap (double positive) objects were exported to Excel (Microsoft Corporation, Redmond, WA) and lethal fraction (LF) scores were computed from mKate2^+^ and SG^+^ counts as described^56^. LF scores were exported to Prism (GraphPad Software, La Jolla, CA) for further analysis.

### Gene expression analysis

For 2D culture, cells were seeded in 6-well plates and harvested at approximately 80% confluency. For 3D culture, 2,000 cells were seeded in 96-well ultra-low attachment (ULA) microplates (Corning, Cat# 7007), spun at 1,200 rpm for 2 min for spheroid formation, then pooled and harvested 48 h post-seeding. Total RNA was extracted using RNeasy Plus Universal Mini Kits (Qiagen, Cat# 73404). For high throughput RT-qPCR, the Biomark/Fluidigm system (Standard BioTools) and TaqMan Gene Expression Assays (Life Technologies) were used. cDNA was pre-amplified with pooled TaqMan Assays and TaqMan PreAmp Master Mix (ThermoFisher, Cat# 4391128). Products were then mixed with pooled TaqMan Gene Expressions Assays, TaqMan Universal PCR Master Mix (Life Technologies, Cat# 4304437) and 20X GE Sample Loading Reagent (Fluidigm, Cat# 100-7610), and loaded onto the 192.24 Dynamic Array integrated fluidic circuit (Fluidigm, Cat# 100-6170) following manufacturer’s instructions. RT-qPCR was conducted as per manufacturer’s protocol. RT-qPCR data was obtained on the microfluidic platform. The mean expression of seven endogenous housekeeping genes across all samples was used for normalization and calculation of -ΔCt values. TaqMan Gene Expression Assays (Applied Biosystems) for targeted genes include: *GPX4* (Hs00157812_m1), *AIFM2/FSP1* (Hs_01097300_m1), *GCH1* (Hs00609198_m1), *DHODH* (Hs00361406_m1), *SLC3A2* (Hs00374243_m1), *SLC7A11* (Hs00921938_m1), *GCLC* (Hs00155249_m1), *GSS* (Hs00609286_m1), *LRP8* (Hs00182998_m1), *NFE2L2* (Hs00975960_m1), *TP53* (Hs01034249_m1), *CDKN1A* (Hs00355782_m1), *ACTB* (Hs99999903_m1), *GAPDH* (Hs99999905)_m1), *HMBS* (Hs00609297_m1), *HPRT1* (Hs02800695_m1), *RPLP0* (Hs99999902_m1), *RPL13A* (Hs01578912_m1) and *TBP* (Hs00427620_m1).

For genome-wide RNA expression, total RNA was extracted (Qiagen, miRNAeasy Mini Kit, Cat# 217084). For sense-strand complementary DNA synthesis, fragmentation and labeling GeneChip^TM^ WT PLUS Reagent Kit (Applied Biosystem, Cat# 902280) was used according to manufacturer’s instructions. DNA target was hybridized on the Clariom D, human array (ThermoFisher, Cat# 902922) and conducted as per manufacturer’s protocol. The Transcriptome Analysis Console software (TAC, Applied Biosystem) was utilized for expression analysis.

### Prostaglandin E_2_ detection

HT-1080 cells were seeded into a 6 well plate. After 24 h, cells were incubated with DuP697 for 72 h. Cell pellets and supernatants were harvested. PGE2 concentration in cell supernatant was measured with Prostaglandin E_2_ ELISA Kit (Cayman) as per manufacturer’s instructions.

### Lipid ROS imaging and quantification

IncuCyte images were acquired on the Sartorius IncuCyte S3 with 10x magnification. For AA and DuP697 experiments, 2,000-4,000 cells were seeded in 96-well flat bottom and ULA plates (Corning, Cat# 7007). AA (10 μM) was added at cell seeding. After 24 h ferrostatin-1 (1 µM) was added to cells 2 h prior to RSL3 treatment. BODIPY 581/591 C11 (Thermo, Cat# D3861) at final concentration 5 mM and DRAQ7 (Novus Bio, Cat# NBP2-81126) at final concentration 3 µM were added to separate wells. RSL3 was added and placed in IncuCyte. The ratio of oxidized to total lipids (C11^Ox^/C11^Non-Ox^ + C11^Ox^) was calculated and graphed with DRAQ7 uptake (Integrated intensity, reference DMSO) at the same time point.

For MUFA experiments, lipid ROS images were acquired on the Lionheart at 20x. For ferrostatin-1 withdrawal experiments 70,000 HT-1080^N^ Control or *GPX4^KO1/2^* cells/well were seeded into 12-well dishes. 24 h later cells were washed three times with 1 mL PBS then were incubated in media with or without ferrostatin-1 (1 μM) for 1.75 h. Medium was aspirated off cells and replaced with 1 mL C11 BODIPY 581/591 (5 μM) and Hoescht (0.1 mg/mL) dissolved in PBS (with 1 μM Fer-1 for the ferrostatin-1 condition) then incubated at 37°C for 15 min. After 15 min, the label was removed and 1 mL of PBS +/- ferrostatin-1 (1 μM) was added to the cells. Cells were imaged immediately using the Lionheart imaging system at 20x magnification, 2 images per well.

For the experiment with 12 h vs 48 h oleic acid (OA) pretreatment, cells were seeded at 3,000 cells per well into a 24-well plate. 24 h after seeding, wells were aspirated, washed with 500 μL PBS, then treated with EtOH or OA (125 μM) to start the 48 h condition, and 36 h later cells were washed with 500 μL PBS before EtOH or OA (125 μM) was applied to wells, starting the 12 h condition. After 12 or 48 h of total EtOH or OA treatment, cells were washed with 500 μL PBS and then regular media was added back to wells for 1 h and 45 min, ferrostatin-1 (1 μM) was only included in media added back to control wells. 500 μL BODIPY 581/591 C11 (5 μM) and Hoescht (0.1 mg/mL) mixture in PBS was applied as described above, and images were acquired on the Lionheart.

For MUFA panel experiments, 2,500 cells were seeded into a 96-well plate. 12 h after cell seeding media was aspirated and 200 μL media containing fatty acids (125 µM) or EtOH vehicle were added to wells. After 12 h of fatty acid treatment, both fatty acids and Fer-1 were withdrawn except for control wells that remained in Fer-1 for 1 h and 45 min. Then, cells were treated with 100 μL C11 BODIPY 581/591 (5 μM) and Hoescht (0.1 mg/mL) as described above and imaged on the Lionheart.

All images for all lipid ROS imaging experiments were captured using the DAPI, Texas Red, and GFP channels were acquired at 20x magnification using the Lionheart instrument. Ratios of red to green fluorescence were calculated in the instrument. Data points were exported into Excel and plotted using Prism. Images were downloaded then processed in FIJI and arranged in Illustrator.

### Neutral lipid staining and quantification

One day prior to imaging, 10,000 HT-1080^N^ GPX4^WT^ cells were seeded into a 96 well plate into media containing ferrostatin-1 (1 μM). Three technical replicates were seeded for each fatty acid treatment condition. 12 h after seeding, cells were treated with fatty acids at 125 μM or EtOH vehicle for 12 h before wells were aspirated and treated with BODIPY 493/503 (1 μM) with ferrostatin-1 (1 μM) in PBS at 37°C for 30 min. After 30 min, cells were washed gently three times with 100 μL PBS. After the final wash, 200 μL of PBS with ferrostatin-1 (1 μM) was added and cells were imaged (40x, 2x2 montage, Lionheart). Three independent biological replicates were performed for each condition. Lipid droplet count from the four total images per technical replicate in each biological replicate was calculated using FIJI.

In other experiments, one day prior to imaging, 10,000 HT-1080^N^ Control or *ACSL3*^KO1/2^ cells per well were seeded into a 96-well plate. 24 h later, media was removed and media containing fatty acids (125 μM) or EtOH control was added to the plate for 12 h. Then, media was removed and treated with 200 μL BODIPY 493/504 (1 μM in PBS) per well for 30 min. After this incubation, BODIPY 493/503-containing medium was removed and replaced with 200 μL PBS before imaging. Lipid droplet counts from four total images per replicate were calculated in FIJI.

### Shotgun lipidomics

For 2D culture versus 3D cell culture experiments, 2D cells were seeded in 10 cm dishes and harvested at approximately 80% confluency. For 3D culture, cells were seeded in 6-well Elplasia plates (Corning, Cat# 4440), spun at 1,200 rpm for 2 m for spheroid formation, then harvested 72 h post-seeding. Cells were washed (PBS 1X, Cat# 10010023, Gibco), harvested (TrypLE, Cat# 12605010, Gibco), counted with ViCell BLU (Beckman Coulter), centrifuged to pellet, then resuspended in PBS. For tumors, approximately 20 mg was harvested. For other studies, 500,000 HT-1080^N^ cells were seeded into 10 cm dishes (Cat# CC7682-3394, USA Scientific, Ocalal, FL). The next day, dishes were treated with fatty acids at 125 μM for 24 h. Then, cells were trypsinized and harvested into media, counted, centrifuged to pellet cells 55 x g, 5 min, room temperature, then washed with PBS. Cells were re-centrifuged then resuspended in a small volume of PBS and frozen at -80°C in flat-bottom, open top glass tubes until lipids were extracted and analyzed as described previously^21^ at the UCLA Lipidomic core. Four biologically independent replicates were performed for each condition.

## QUANTIFICATION AND STATISTICAL ANALYSIS

Lethal fraction scoring was performed using Microsoft Excel 14.6.0 (Microsoft Corporation, Redmond, WA) and Prism 9 or 10 (GraphPad Software, La Jolla, CA). Graphing and statistical analyses were performed using Prism 9 or 10. Figures were assembled using Adobe Illustrator (Adobe Systems, San Jose, CA). Statistical details of experiments and statistical tests used can be found in the main text and figure legends.

## DATA AND CODE AVAILABILITY

The data generated in this study are publicly available in Gene Expression Omnibus (GEO) and Mendeley.

---

**Figure S1.**
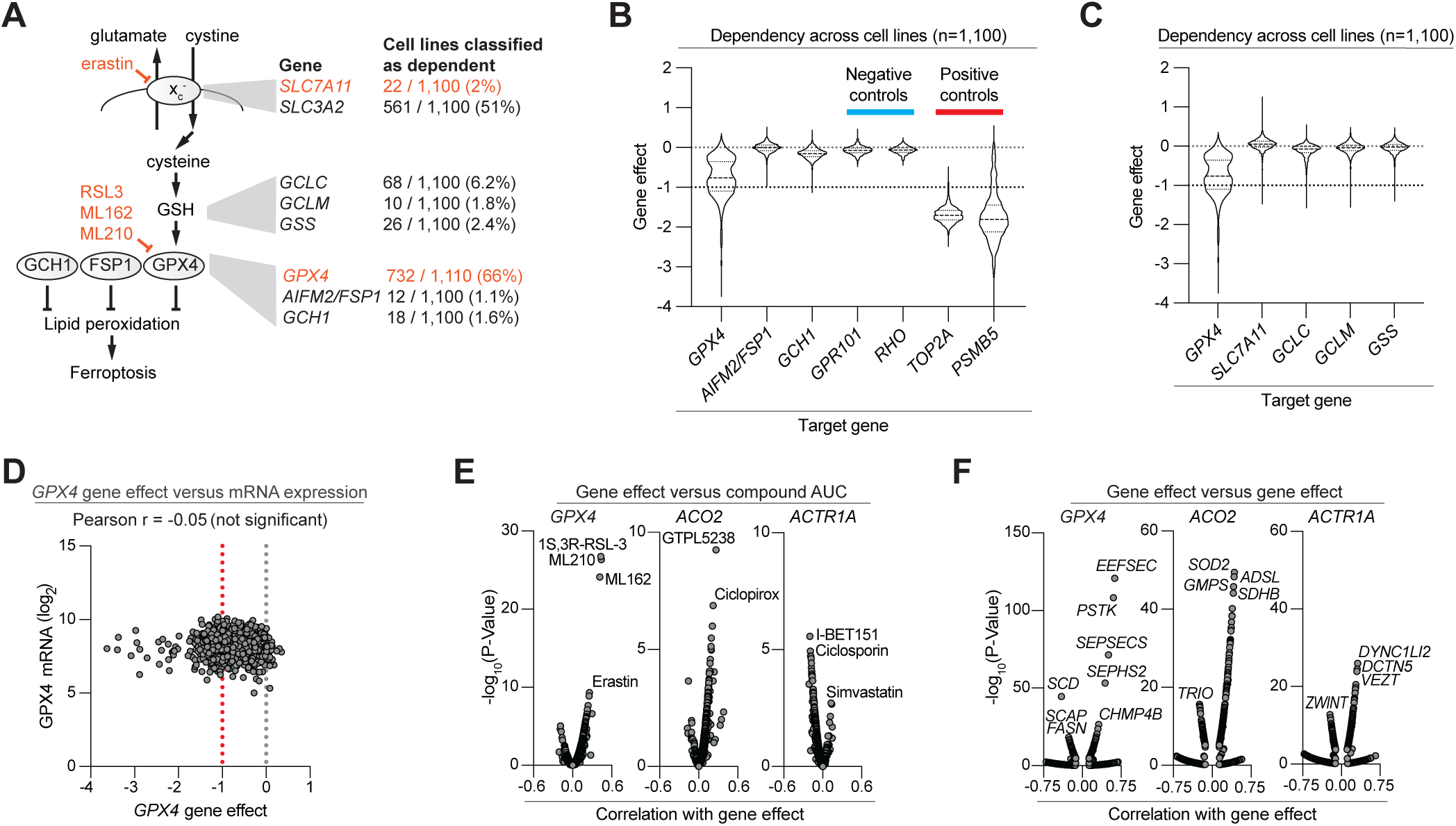
Analysis of ferroptosis pathway genes using the DepMap resource. **A**, Cartoon summary of the ferroptosis mechanism with key anti-ferroptotic genes highlighted. On the right, a summary of results from DepMap (24Q2 release) with the number of cell lines classified as being dependent on a given gene. **B**, Data extracted from DepMap showing gene effect scores accross all 1,100 cell lines. *GPR101* and *RHO* are control non-essential genes, while *TOP2A* and *PSMB5* are positive control essential genes. **C**, Extracted gene effect data for *GPX4* compared to genes in the upstream cystine and GSH metabolic pathway. **D**, *GPX4* gene effect scores versus mRNA expression. Each datapoint represents a single cell line. **E**, Correlations of gene effect scores with small molecule effects. *ACO2* and *ACTR1A* were identified as control genes that, like *GPX4*, are dependencies in ∼66% of all cancer cell lines in DepMap. **F**, Correlation of gene effect scores with other gene effect scores.

**Figure S2.**
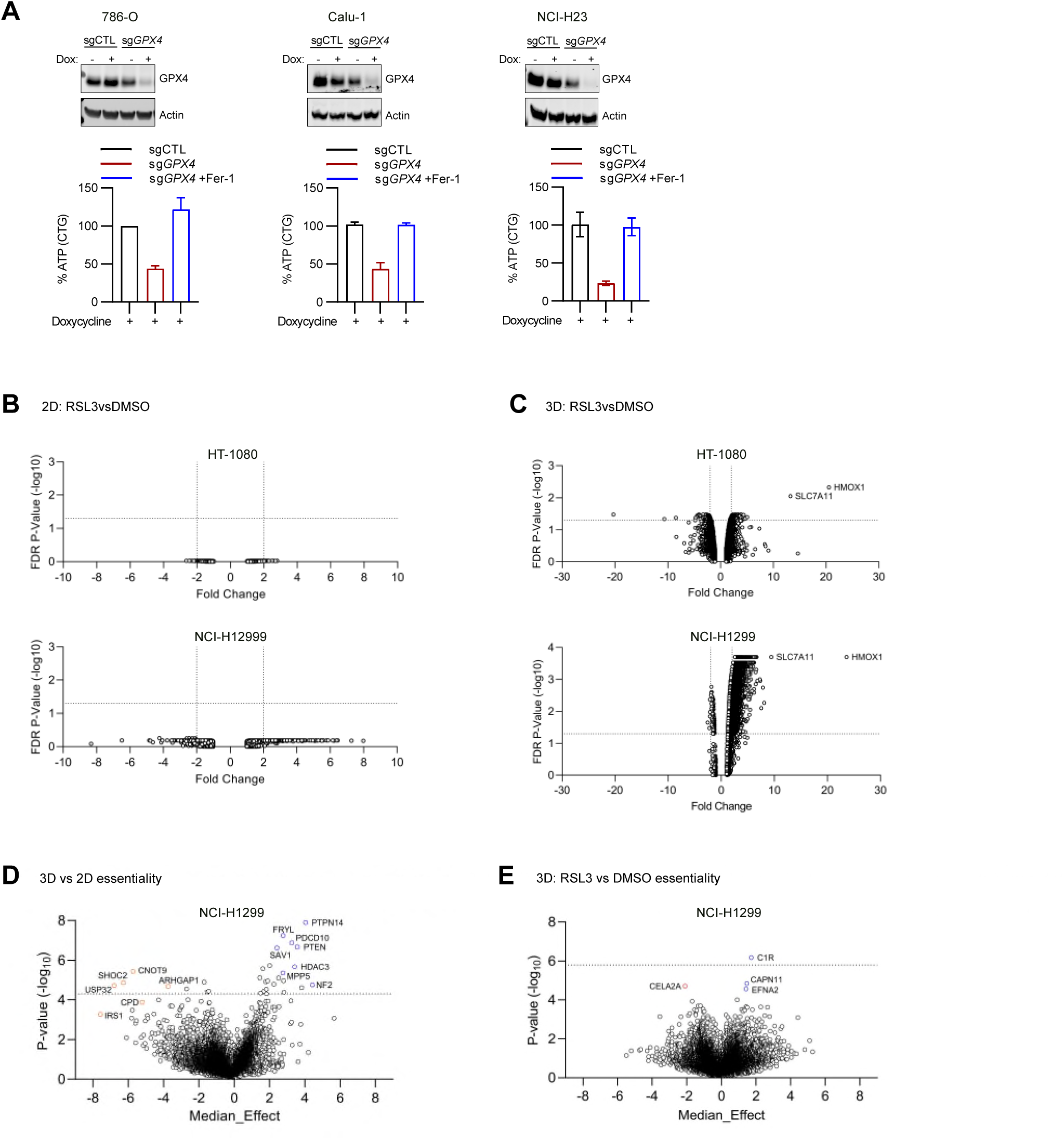
Experimental systems affect transcriptional changes when ferroptosis is induced. **A.** Inducible GPX4 western blot. **B-C.** Volcano plots of genome-wide differential expression measured via microarray (Clariom D assay). **B.** Differential expression of 2D cells 2 hours post-RSL3 (0.1 μM) treatment. Median of 4 biological replicates is represented, FDR *P* <0.05. **C.** Differential expression of 3D cells 24 hours post-RSL3 (0.1 μM) treatment. Median of 4 biological replicates is represented, FDR *P* <0.05. **D.** 3D genome-wide CRISPR screen in NCI-H1299 enCas12a cells comparing 3D culture vs 2D culture depletion and enrichment of gene disruptions. **E.** 3D genome-wide CRISPR screen in NCI-H1299 enCas12a cells comparing 3D RSL3 (2 μM) vs 3D DMSO depletion and enrichment of gene disruptions. Analyses are based on 4 biological replicates for each condition. Hashed line delineates FDR *P* <0.05.

**Figure S3.**
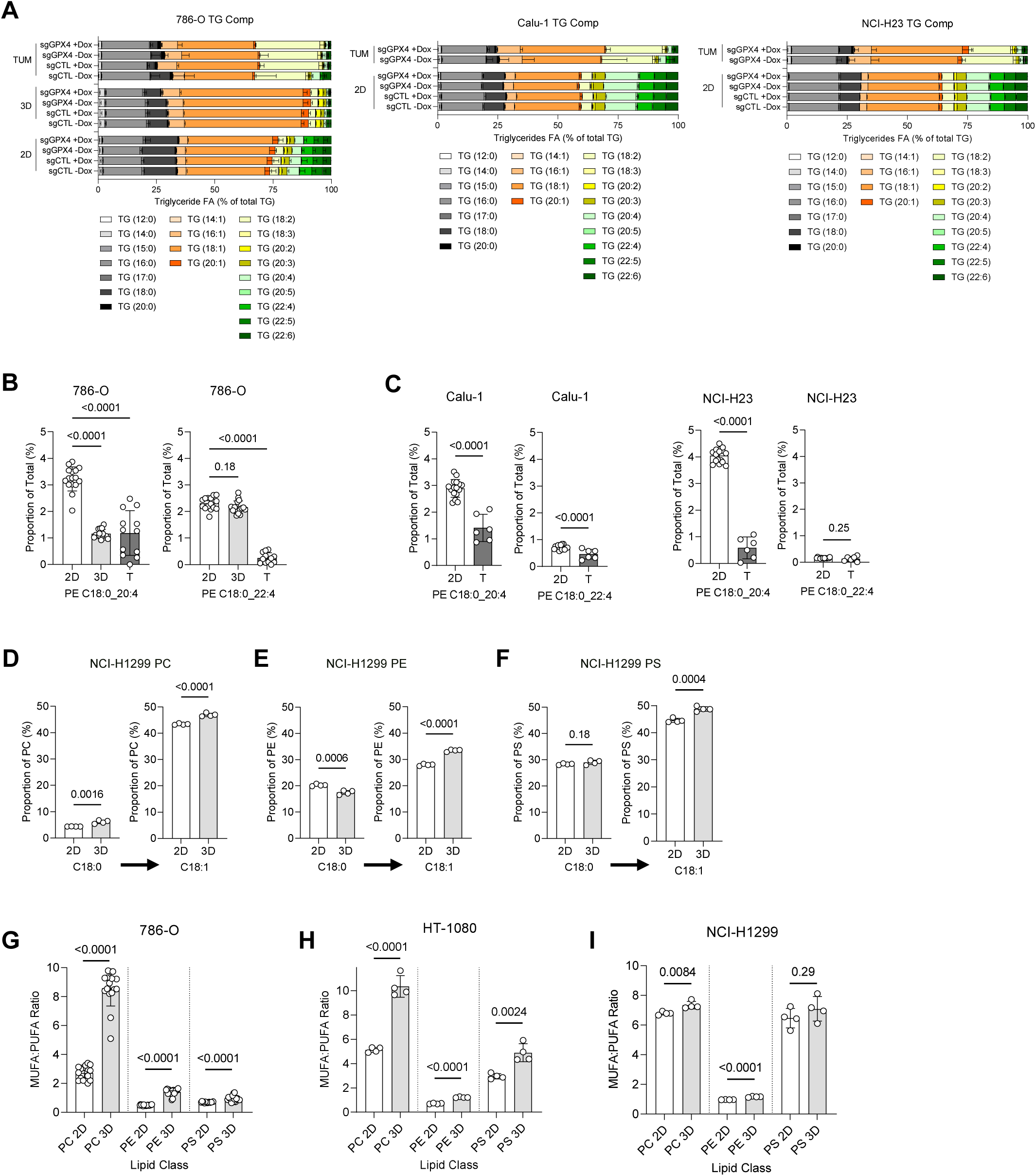
Experimental systems affect transcriptional changes when ferroptosis is induced. **A.** Proportion of fatty acids (FAs) within TGs in 2D cells, 3D cells and tumors in 786-O cell line. FAs are categorized by the number of carbons and double bonds. Stacked bar graphs are presented as means ± SD. **B-C.** Proportion of PE 18:0_20:4 (AA) compared to the total lipidome in 2D cells, 3D cells and tumors in 786-O, Calu-1 and NCI-H23 cell lines. **D-F.** Changes in 18:0-PL (saturated fatty acid, SFA) ◊ 18:1-PL within PC (**D**), PE (**E**) and PS (**F**) in 2D cells and 3D cells of NCI-H1299 cell line. In B-F, data are presented as means ± SD and statistical significance was assessed using two-tailed unpaired Student’s t-test. **G-I.** Ratio of proportion of all MUFAs to all PUFAs in PC, PE and PS in 786-O, HT-1080 and NCI-1299 cell lines.

**Figure S4.**
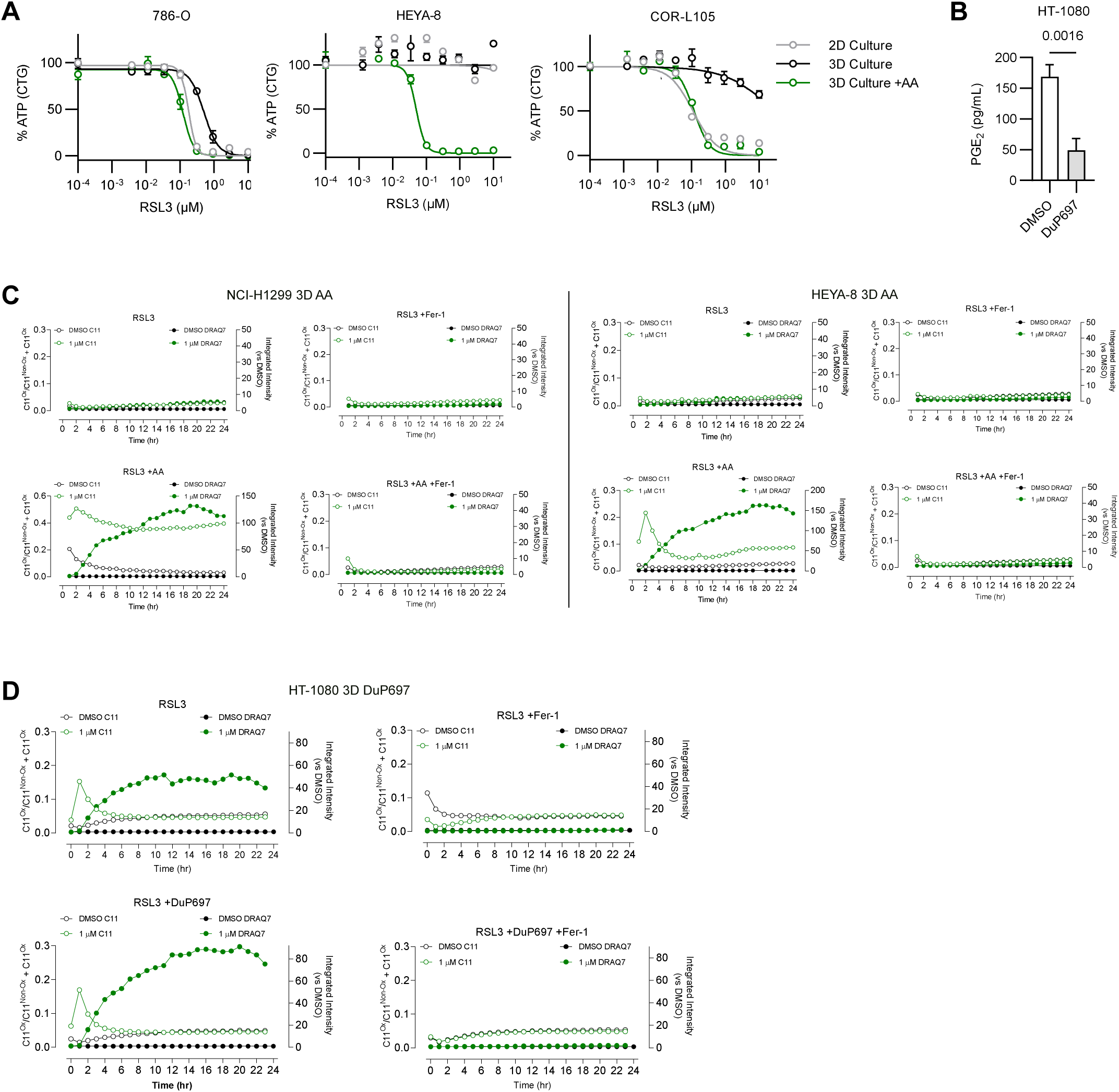
Increasing arachidonic acid sensitizes cells to ferroptosis induction in 3D cell culture. **A.** ATP levels (CTG) for 2D and 3D cultures of COR-L105, HEYA-8 and 786-O cells, with AA (10 μM) added to cell media at seeding, then treated with RSL3. **B.** PGE_2_ levels in 2D and 3D cells following 48 h DuP697 (10 μM) treatment. **C.** Lipid peroxidation (BODIPY 581/591 C11) and cell death (DRAQ7) of NCI-H1299 and HEYA-8 3D cells pre-treated with AA (10 μM), RSL3 and ferrostatin-1 (1 μM). **D.** Lipid peroxidation (BODIPY 581/591 C11) and cell death (DRAQ7) of HT-1080 3D cells pre-treated with COX2 inhibitor (DuP697, 10 μM) and at seeding, then treated with RSL3. Data represent biological replicates.

**Figure S5.**
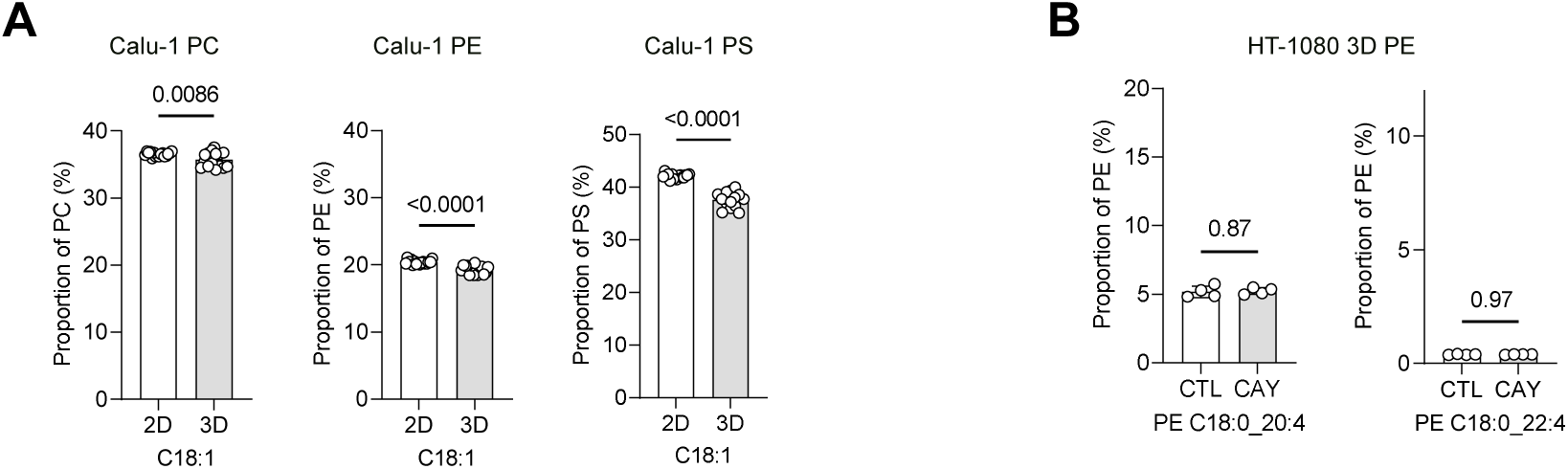
Decreasing SCD products sensitizes cells to ferroptosis induction in 3D cell culture. **A.** Proportion of C18:1-PL (OA) within PC, PE and PS in 2D cells and 3D cells of Calu-1 cell line. **B.** Proportion of lipid species PE-AA and PE-AdA within PEs in HT-1080 3D cells comparing control (2% media) and SCD1 inhibitor (CAY10566, 200 nM) overnight. Data are presented as means ± SD and statistical significance was assessed using two-tailed unpaired Student’s t-test.

**Figure S6.**
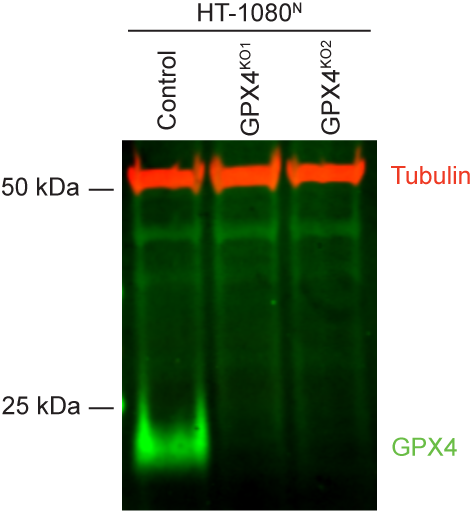
GPX4 protein expression. Immunoblot of HT-1080^N^ Control and *GPX4^KO1/2^* cells. Note that prior to cell lysis *GPX4^KO1/2^* cells were cultured with ferrostatin-1 (1 µM). Representative of two experiments.

**Figure S7.**
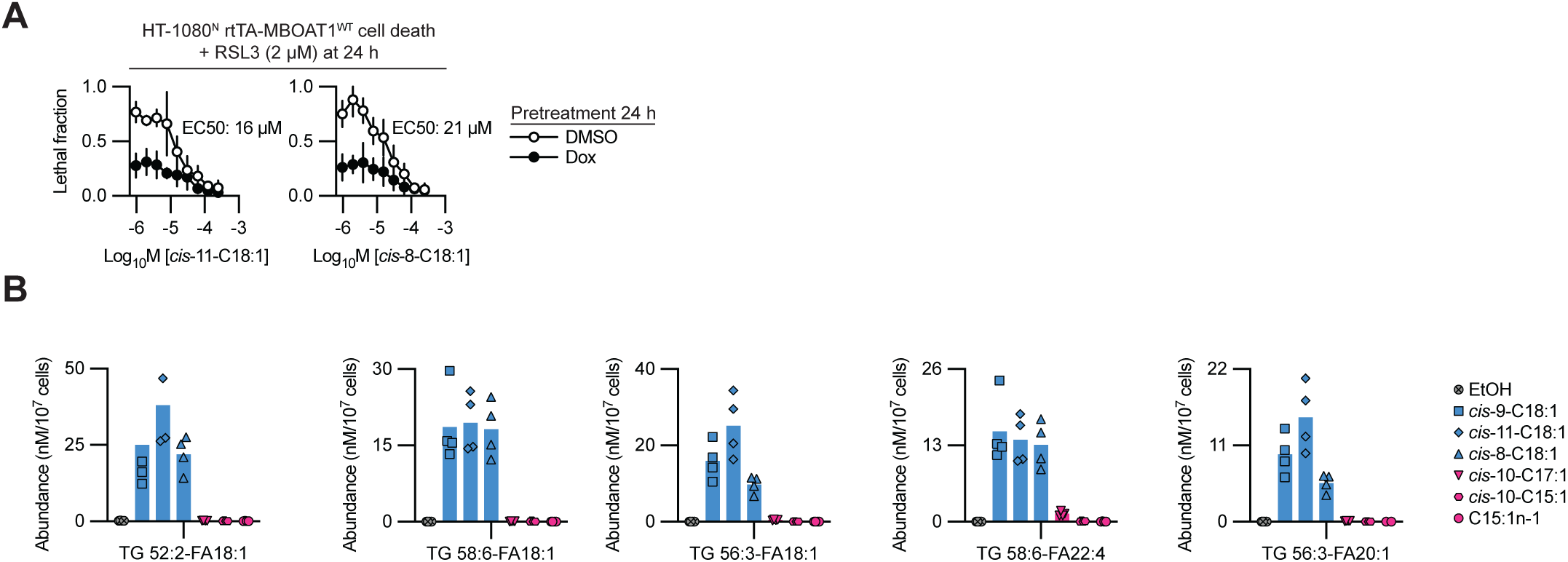
MUFAs impact ferroptosis sensitivity. **A**, Cell death determined by imaging. Cells were pretreated for 24 h ± doxycycline (Dox, 1 µg/mL) and various concentrations of MUFAs, prior to RSL3 treatment. Results are mean ± SD from three independent experiments. **B**, Lipid abundance determined by shotgun lipidomics. Four independent samples were collected per condition.

